# Living xylem cells encode a large number of conserved gene families responsible for vascular sap peptides

**DOI:** 10.1101/2024.11.10.622827

**Authors:** Min R. Lu, Chia-Chun Tung, Chia-En Huang, Chang-Hung Chen, Pin-Chien Liou, Chan-Yi Ivy Lin, Jhong-He Yu, Ying-Lan Chen, Ying-Chung Jimmy Lin, Isheng Jason Tsai

## Abstract

Plant long-distance signaling peptides travel through the vascular system to coordinate development and respond to environmental cues, yet their precursor genes and expression origins remain elusive. We characterized 4,804 sap peptide precursor genes in *Populus trichocarpa* using an integrated approach that combined liquid chromatography tandem mass spectrometry (LC-MS/MS) peptidomics, transcriptomics, and comparative genomics. This study expands the known precursor families from approximately 50 to thousands, the majority of which are conserved across angiosperms. Transcriptome analysis across xylem developmental stages revealed that living xylem cells, typically viewed as precursors to non-living structures, predominantly express these specifically at transitions between primary and secondary growth stages, indicating an active role in plant-wide signaling coordination. The precursor genes show conservation at the transcriptome level and are under strong purifying selection. Our findings provide a comprehensive overview of the gene families encoding sap peptides, redefining xylem as an active participant in plant communication and adaptation.

## Introduction

Signaling peptides play a pivotal role in plant development and environmental response, functioning analogously to hormones by regulating various physiological processes^1–3^. These peptides, once recognized by cell membrane receptors, initiate intracellular signaling cascades that modulate downstream responses. Known signaling peptides are derived from small secreted proteins (SSPs), typically ranging from 100 to 250 amino acids^4,5^, which are processed by proteases into peptide cleavage products^6,7^ (PCPs) and synthesized in specific cells or tissues before being transported to their target sites. Based on their mode of action, they can be classified as either short-distance peptides, which mediate cell-cell communication, or long-distance peptides, which facilitate signaling between different organs and tissues^8–12^. Understanding the functional roles of these peptides is critical for uncovering the mechanisms of long-distance communication in plants, a field that is still emerging^13–15^. Another key question is the diversity and function of the precursor genes encoding these peptides. While *in silico* approaches predict thousands of such genes across species, fewer than a hundred precursor gene families have been experimentally validated in each species to date^16,17^.

While the functions and origins of short-distance peptides have been extensively characterized—playing critical roles in apical and lateral growth^3,18– 31^ —our understanding of long-distance peptides remains limited, particularly with respect to their production, mechanisms, and functional roles in plant biology. Vascular sap serves as a crucial conduit for water, nutrients, and signaling molecules including small peptides. The known long-distance mobile peptides modulate various physiological processes, including water deficiency responses and pathogen defense. For example, peptide such as CLAVATA3/EMBRYO-SURROUNDING REGION-RELATED 25 (CLE25) have been identified as mobile signals which is produced in roots and travels through the vasculature to the shoot, where it influences stomatal behavior to mitigate water loss under drought conditions^31^. Recent advances, including liquid chromatography tandem mass spectrometry (LC-MS/MS) and peptidomic profiling methods^32^, have allowed the identification of PCPs systematically, some of which were identified in vascular sap^10,33,34^. These sap peptides appear to be conserved across angiosperms, and new insights are emerging regarding their translocation and functional roles within plants^34^. The growing body of evidence points to their significance in long-distance signaling, highlighting the need to uncover the precursor genes that encode them and their possible origins.

Xylem, the most abundant tissue on Earth, plays a critical role in plant biology through its well-known functions of water transportation and mechanical support. Vessel elements and libriform fibers, the two major cell types in xylem, are primarily responsible for these transport and structural roles, facilitated by the heavy deposition of secondary cell walls (SCW) followed by programmed cell death^35^. Several lines of evidences have pointed that living xylem cells could serve as a previously overlooked source of signaling peptide precursor genes^10,31^. For example, genes encoding Xylem sap-Associated Peptides (XAPs) were found to express in the first internode of soybean *Glycine max*^10^, indicating that their expression is physically linked to the vascular system. *Populus trichocarpa*, a model organism for studying wood formation and vascular development^19,36–38^, offers an ideal system for this research due to its compact, well-annotated genome and suitability for omics approaches. As a woody species, *P. trichocarpa* provides a unique opportunity to explore the role of signaling peptides in xylem development, offering insights into broader mechanisms of vascular signaling and tree growth.

We hypothesize that the sap peptide precursor genes are expressed at specific stages of xylem development and are subsequently processed into bioactive peptides, integrating into the plant’s long-distance signaling network. To test this hypothesis, we focused on *Populus trichocarpa*, a well-annotated woody species ideal for studying xylem formation and vascular signaling, as well as two other woody species *Eucalyptus grandis* and *Cinnamomum kanehirae*. We collected sap samples and used LC-MS/MS for peptide identification alongside RNA sequencing for transcriptomic analysis. After identifying sap peptide precursors, we examined their orthology and expression patterns across various developmental stages and representative plant species to investigate conservation and diversity. This integrative approach allowed us to uncover the molecular mechanisms behind initial stages long-distance peptide signaling, providing a deeper understanding of the active roles that living xylem cells play in plant communication networks.

## Results

### A large number of sap peptide precursor gene families conserved across angiosperms

To investigate the diversity of signaling peptides in the vascular sap of woody plants, we constructed a comprehensive peptidomic dataset for *Populus trichocarpa* using an LC-MS/MS-based peptidomic approach **(Figure 1a and Methods)**. This analysis identified 7,039 vascular sap peptides, with an average length of 15 amino acid residues (**Supplementary Figure 1a and Supplementary Table 1**). Mapping these peptides to the *P. trichocarpa* proteome annotation revealed that they originated from 2,254 unique precursor proteins **(Figure 1b and Supplementary Table 2)**. The non-redundant matched peptide regions typically spanned 11 to 16 amino acids, while the majority of these precursor proteins were between 315 and 550 amino acids **(Figure 1c)**. Only 10.3% of these precursor genes contained a canonical N-terminal signal sequence. Orthogroup inference across 14 representative species **(Supplementary Table 3)** placed the *P. trichocarpa* genes with matched sap peptides into 1,374 orthogroups (**Supplementary Table 4**). Random sampling of an equivalent number of genes without replacement yielded an average of 2,034 orthogroups (**Supplementary Figure 2**), indicating that precursor genes were members of gene families rather than single copy genes—a result that significantly deviated from chance (P < 0.01, Bootstrapping, **Supplementary Figure 2**). We observed that peptide-matched regions were not fully conserved between orthologs, indicating variable sequence conservation among sap peptides **(Figure 1b)**. Nevertheless, the rarefaction curve for these orthogroups approached a plateau **(Figure 1d)**, and 44.1% (606/1,374) of the orthogroups contained all gene members matched to sap peptides (**Supplementary Figure 3**), suggesting that our peptidomic approach effectively captured the majority of the precursor gene families.

**Fig. 1.**
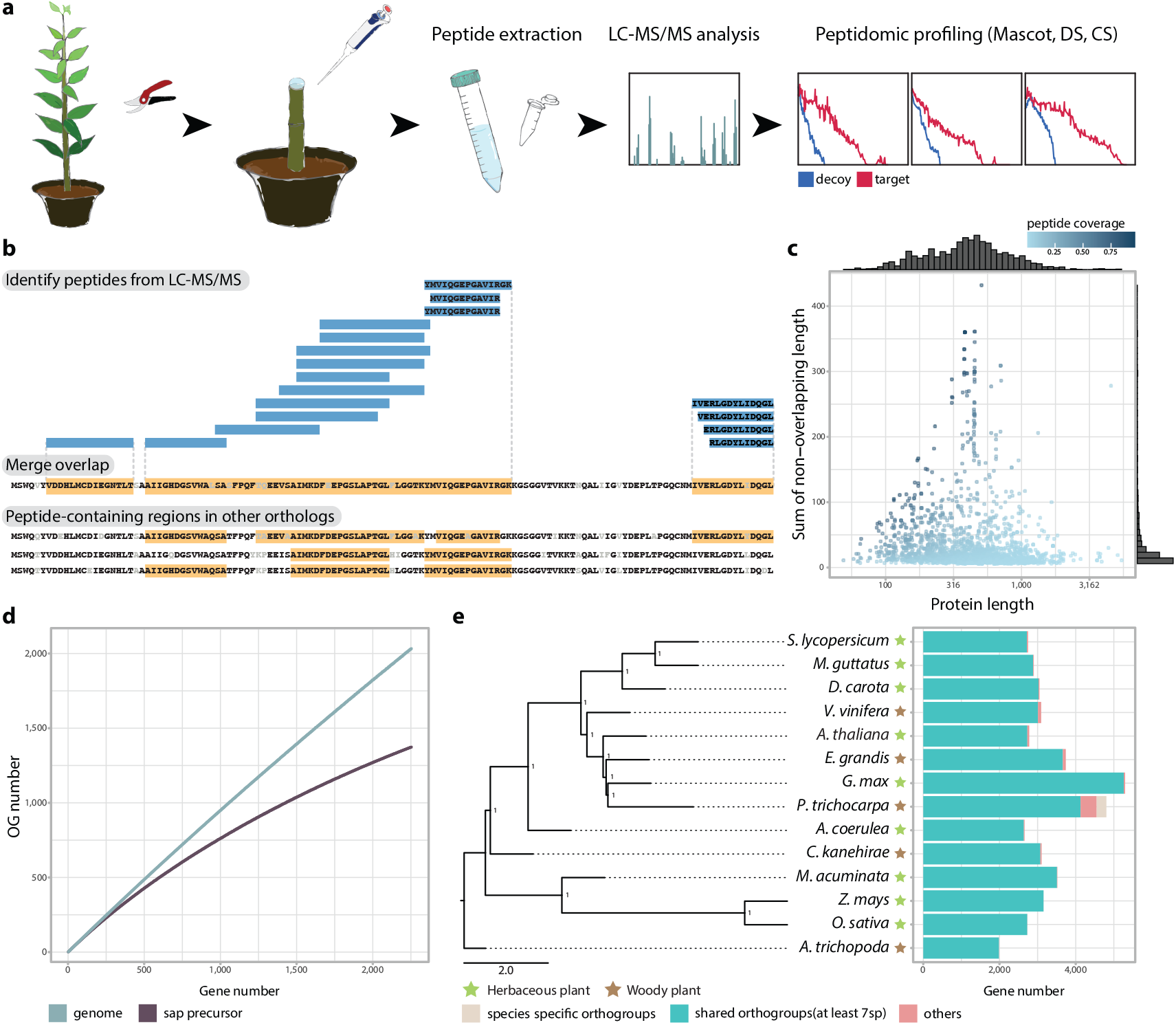
Characterization of vascular sap peptides in *Populus trichocarpa*. (a) Schematic of the experimental workflow for vascular sap peptide extraction and identification using an LC-MS/MS-based peptidomic profiling approach (**Methods**) (b) Schematic of identifying sap peptide precursors. Sap peptides were identified across putative precursors using endogenous peptide identification pipeline^32^. Overlapping regions of sap peptides within proteins were merged. The figure shows different peptide-matching patterns within the same orthogroup OG0000404. (c) Distribution of sap precursor protein length and sum of regions with evidence of peptide sequence matching. Color of points denote the proportion of peptide regions within genes. (d) Orthogroup analysis from random sampling of 1 to 2,254 genes, with each point representing the average orthogroup count from 100 iterations. Colors distinguish whether the selection is from the entire genome or specifically from sap peptide precursors. (e) Phylogenetic tree of 14 representative plant species, with herbaceous and woody species indicated by color. The right panel shows the distribution of *P. trichocarpa* sap peptide precursor orthologs across species, highlighting genes shared by at least half of the species and those unique to *P. trichocarpa*.

We designated a total of 4,804 *P. trichocarpa* genes from these orthogroups as sap peptide precursor genes and found that 91.1% of them have orthologs in at least half of the representative species, suggesting strong conservation across angiosperms **(Figure 1e)**. Known gene families among these precursors include tubulin and ubiquitin (**Supplementary Table 5**), consistent with previous findings that plant peptidomes contain proteolytic cleavage products (PCPs) from the degradation of functional proteins^33,39^. In addition, only 45% of these *P. trichocarpa* genes were annotated with gene ontology (GO) terms. These precursor genes were not enriched for any specific biological processes, indicating that their primary role is in the generation of sap peptides. Interestingly, the proportion of disordered regions was higher in sap peptide precursor genes without N-terminal signal sequence predicted, compared to both non-precursor genes and those with signal sequence (Wilcoxon rank sum test, P < 0.05, **Supplementary Figure 4a**). This proportion also correlated with the extent of peptide-matched regions (Spearman correlation, **Supplementary Figure 4b**), suggesting that disordered regions might may facilitate structural flexibility, allowing the peptides to adopt conformations necessary for biological activity or interaction with receptors.

To determine if the identified *P. trichocarpa* precursor genes were consistent with existing datasets, we reanalyzed peptide PCPs previously identified in hybrid poplar *Populus x canescens* during interaction with the ectomycorrhizal basidiomycete *Laccaria bicolor*^*33*^. We found that 17 out of 25 xylem sap specific precursor orthogroups defined in REF33 were also designated as precursors in our dataset, a significantly enrichment compared to non-xylem precursors (X^2^ test, P=0.006075, **Supplementary Table 6**), suggesting that our precursor datasets were robust and consistent with known xylem sap precursors. Next, we compared the *P. trichocarpa* precursor genes to their orthologs in *Arabidopsis thaliana* and *Medicago truncatula*, both of which have databases containing limited numbers of experimentally validated and *in silico*-predicted small secreted peptides (SSPs)^16,17^. Although more than half of the SSPs in these species shared orthology with *P. trichocarpa* genes, only a small fraction showed matching peptide evidence. For instance, of the 1,023 SSPs matched to 764 genes in *A. thaliana*, 185 orthogroups were identified, with 100 shared with *P. trichocarpa*; however, six of these orthogroups contained *P. trichocarpa* sap peptide precursor genes, indicating limited overlap and differences in peptidomes **(Supplementary Figure 5)**. In contrast, among the 231 long-distance mobile peptides identified in soybean (*Glycine max*) xylem sap^10^, 37 orthogroups were identified, with 35 shared with *P. trichocarpa*. Of these orthogroups, 45.7% included 61 *P. trichocarpa* sap peptide precursor genes **(Supplementary Figure 5)**. Further analysis of 12 *Glycine max* Xylem sap-Associated Peptides (XAPs) predicted as functional^10^ and expressed in various organs, including the xylem sap collection site, showed that six shared *P. trichocarpa* precursor orthologs with two of them matching peptides (**Supplementary Table 7**). The peptides were similar and consistently mapped to corresponding regions within the precursor genes across both species **(Supplementary Figure 6)**, indicating structural conservation and functional relevance. These results highlight the complexity and evolutionary divergence of sap peptidomes while also suggesting potential conserved roles for xylem-derived peptides in long-distance signaling.

### Elevated expression of sap precursor genes in xylem-associated tissues

Transcriptomic analysis across multiple tissues of *P. trichocarpa* revealed that sap precursor genes collectively exhibited a higher proportion of expression in xylem-associated tissues (34.8% to 39.9% versus others 20.9% to 31.4%; **Figure 2a, Supplementary Table 8**). We also generated tissue transcriptomes for *Cinnamomum kanehirae*, and found that this expression ratio was consistent in the expression of 3,105-3,739 sap precursor orthologs in the woody species *E. grandis* and *C. kanehirae*. In contrast, the herbaceous plant *A. thaliana* showed a divergent expression pattern, likely due to differences in xylem cell composition and their development, such as the absence of the ray parenchyma cells and the immature development of libriform fibers in *A. thaliana* (**Figure 2a**). Nevertheless, the highest expression levels of sap precursor orthologs in *A. thaliana* were found in vascular bundles, suggesting that sap precursor genes have diverse and potentially important roles within xylem tissues across a broad range of species.

**Fig. 2.**
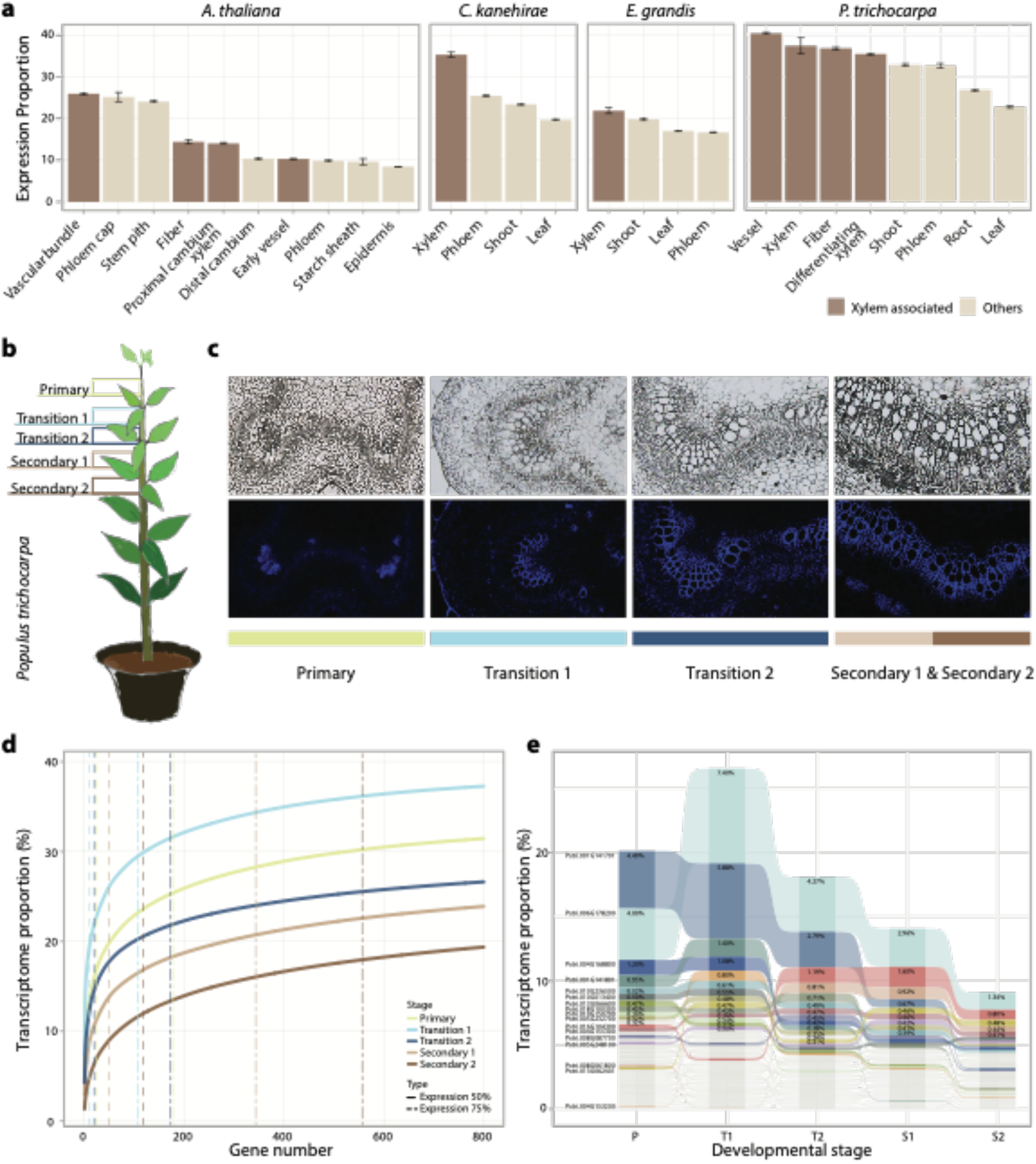
Xylem specific expression of sap peptide precursor genes across angiosperms. (a) Proportion of sap peptide precursor gene expression across various tissues in different plant species, with color coding indicating whether the tissue is associated with xylem. (b) Schematic of *Populus trichocarpa* illustrating the relative positions of the five developmental stages analyzed. (c) Cross-sections of stem tissue during wood formation, highlighting the blue-stained living xylem cells selected for laser-capture microdissection (LCM). (d) Cumulative expression proportion of sap peptide precursor genes across developmental stages in *P. trichocarpa*. Dashed lines indicate the number of genes contributing to 50% and 75% of total expression, ranked from highest to lowest expression proportion. Note that gene ranking varies by developmental stage. (e) Expression profiles of sap peptide precursor genes during different developmental stages in *P. trichocarpa*. Only genes contributing more than 0.1% to the total transcriptome are shown. Genes are ranked by expression levels, with the top ten genes at each stage highlighted for clarity.

### Expression of sap peptide precursor genes dominated transcriptome of early xylem development

The consistent dominance of sap peptide precursor genes in the xylem transcriptome suggests their involvement xylem development. To determine their source, we used laser capture microdissection (LCM) to further partition xylem development into distinct stages. Anatomical analyses of stem cross-section in *P. trichocarpa* and another woody species *E. grandis* allowed us to identify stem internodes representing late primary and early secondary growth stages (**Figure 2b and Supplementary Figure 7**). During primary growth, the vasculature was organized in bundles, forming an eustele architecture **(Figure 2c)**. As the plant transitioned to secondary growth, the vascular tissue reorganized into a circular pattern, with the emergence of libriform fibers as key morphological indicator **(Figure 2c)**. These libriform fibers, characterized by thickened secondary cell walls, appeared during secondary growth to provide mechanical support, but were absent during primary growth. We designated these stages as primary growth (P), two transition stages (T1 and T2) and the two secondary growth stages (S1 and S2) and profiled their corresponding xylem transcriptomes. Sap peptide precursor genes contributed significantly to the transcriptome, ranging from 23.9% to 39.3% across developmental stages (**Figure 2d and Supplementary Figure 8**). The highest expression of sap peptide precursor genes occurred during the first transition stage (T1), followed by adjacent stages, suggesting that sap peptides originate from precursor genes expressed early in axial xylem development. Among the precursor genes, a small subset dominated the transcriptomes; for example, Potri.006G178200 and Potri.001G141701, both annotated as hypothetical proteins, accounted for 7.4% and 5.9% of the T1 transcriptome, respectively. Overall, the top 25, 11 and 20 expressed genes accounted for half of the precursor gene expression in P, T1 and T2 stages, respectively. Approximately half of the sap precursor gene orthogroups were in the top 25% highest expression category in *P. trichocarpa* (X^2^ test, P < 0.01), which included members of XAP gene family **(Supplementary Figure 9)**. Variations in expression patterns between stages were observed **(Figure 2e)**, suggesting dynamic regulation of sap precursor genes throughout xylem development. These highly expressed genes ranged from 100 to 450 amino acids in length and contained up to five matched peptide regions (**Supplementary Figure 10**). In addition, their expression trend appeared to be grouped according to the relative position of the peptides within the precursor genes (**Supplementary Figure 11**).

To investigate whether the orthologs of *P. trichocarpa* sap peptide precursor genes exhibit high expression levels across xylem developmental stages in another woody species, we compared the corresponding transcriptomes in *E. grandis*. (**Supplementary Figure 12**). In *E. grandis*, 3,739 orthologs accounted for 20.5% to 25.4% of the transcriptome across the same designated xylem developmental stages, with collective expression levels peaking during the T1 stage **(Supplementary Figure 12a**). To validate whether these orthologs were indeed sources of sap peptides, we re-analyzed the *E. grandis* sap peptidome^34^ and mapped 331 identified peptides to 505 genes. Of these, 47.5% shared orthology with *P. trichocarpa* precursor genes, while 4.8% had no orthologs in other species. The combined dataset now accounts for 25.4% to 28.8% of the transcriptome across the same five xylem development stages, with expression consistently peaking during the T1 stage **(Supplementary Figure 12a and 12b**). This pattern suggests a common evolutionary origin for sap peptides in woody species. We also observed that a few genes dominated the xylem transcriptome in *E. grandis*^40^ **(Supplementary Figure 13)**; however, expression trends differed between the two species. In *P. trichocarpa*, the majority of highly-expressed precursor genes peaked during the T1 or P stages, followed by a decline in later stages. In contrast, highly-expressed precursor genes in *E. grandis* either maintained consistent high or showed increased expression towards in later developmental stages. For example, Eucgr.K00671 showed more than twice the expression in the S2 stage compared to the P and T1 stages. These results indicate complex regulatory mechanisms and diverse functions of sap peptide precursor genes across species and developmental stages.

### The sap peptide precursor gene families are evolutionarily conserved across angiosperm species

To determine if the expression of sap peptide precursor genes is conserved beyond eudicots, we analyzed transcriptome correlations between the five xylem stages of *P. trichocarpa* and different tissues in representative species (**Figure 3a**). Our analysis revealed that the xylem stages of *P. trichocarpa* displayed higher correlation with xylem-associated stages in other species, with correlation values reflecting the evolutionary divergence between species. In addition, precursor genes and their orthologs showed significantly higher correlations (P < 0.01, bootstrapping; **Supplementary Figure 14** and **15**), suggesting that these genes are more evolutionarily conserved than others. Although no clear clustering was observed between xylem developmental stages of *P. trichocarpa* and *E. grandis* (**Figure 3a**), precursor genes remained more correlated and constrained (**Methods**) compared to other orthogroups, emphasising their conservation at the transcriptome level. The average ratio of nonsynonymous to synonymous substitution rates (*d*N/*d*S) were significantly lower in precursor genes compared to the rest of the proteomes (*P. trichocarpa*-*E. grandis*, P = 9.17e-18; *P. trichocarpa*-*C. kanehirae*, P = 2.50e-19), suggesting stronger purifying selection **(Figure 3b)**. Within the coding regions of these genes, sequence identity was also higher in the peptide-containing regions **(Supplementary Figure 16)**. Despite the conservation, when examining the top ten expressed orthogroups, only half were shared between *E. grandis* and *C. kanehirae* (**Figure 3c and Supplementary Figure 17**), while other pairwise species comparisons showed a dominance of species-specific gene families. These results suggest that although the sap peptide precursor genes were evolutionarily conserved and subjected to strong negative selection, species-specific adaptations also occurred, likely reflecting the unique physiological needs of different species.

**Fig. 3.**
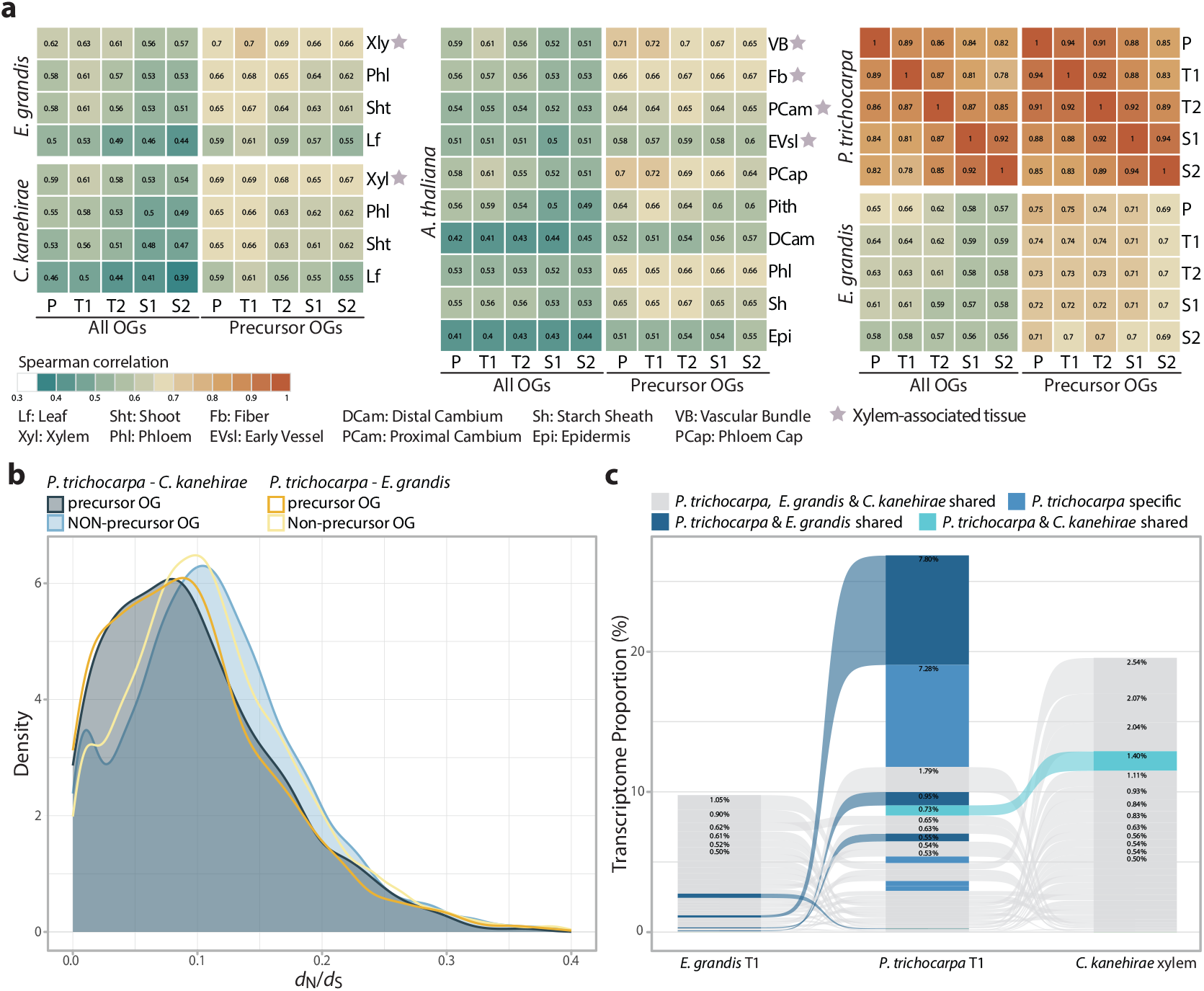
Conservation of gene of sap peptide precursor genes. *(a)* Correlation matrix displaying gene expression patterns across different tissues and developmental stages in eudicots. Correlations are presented for all orthogroups (left) and specifically for those containing sap peptide precursor genes (right). Tissues associated with xylem are marked with stars. *(b)* Density plot of *d*N/*d*S values for orthologs between *P. trichocarpa* and *E. grandis* as well as *P. trichocarpa* and *C. kanehirae*. Colors differentiate precursor from non-precursor orthologous sequences. The plot includes only orthologs with *d*N/*d*S values below 0.4, highlighting the stronger purifying selection on precursor genes. In total, 11,903 orthologs were identified between *P. trichocarpa* and *E. grandis*, and 11,410 between *P. trichocarpa* and *C. kanehirae*, with 1,223 and 1,201 from precursor gene families, respectively. *(c)* Comparative expression of orthologous sap peptide precursor genes across *P. trichocarpa, E. grandis*, and *C. kanehirae*. Only orthologs with expression levels above 0.3% of the respective transcriptomes are shown, with colors representing the orthology relationships among the species.

To validate the conservation of sap peptide precursor genes at the peptide sequence level, we characterised the peptidome of another woody species, *C. kanehirae*, and reanalyzed the peptidomes of both *P. trichocarpa* and *E. grandis*. Given the fewer identified peptides in *C. kanehirae* and *E. grandis* compared to *P. trichocarpa*, only 45 orthogroups containing peptide precursor genes were consistently identified across all three species, whereas 157 orthogroups were shared between *P. trichocarpa* and one of the other species **(Supplementary Figure 18)**. Despite these differences, both shared and distinct peptide features were observed between orthologs within and across species (**Supplementary Figure 19**). For example, in orthogroup OG0000384 **(Figure 4a)**, three peptide regions were uniquely identified in *P. trichocarpa*, while the remaining nine peptide regions were shared across all three species, albeit with slight variations in peptide boundaries. In addition, we found an enrichment of lysine residues at putative peptide cleavage sites in both *P. trichocarpa* and *C. kanehirae* **(Figure 4b)**, suggesting the possible involvement of a homologous protease responsible for cleaving sap peptides in these species.

**Fig. 4.**
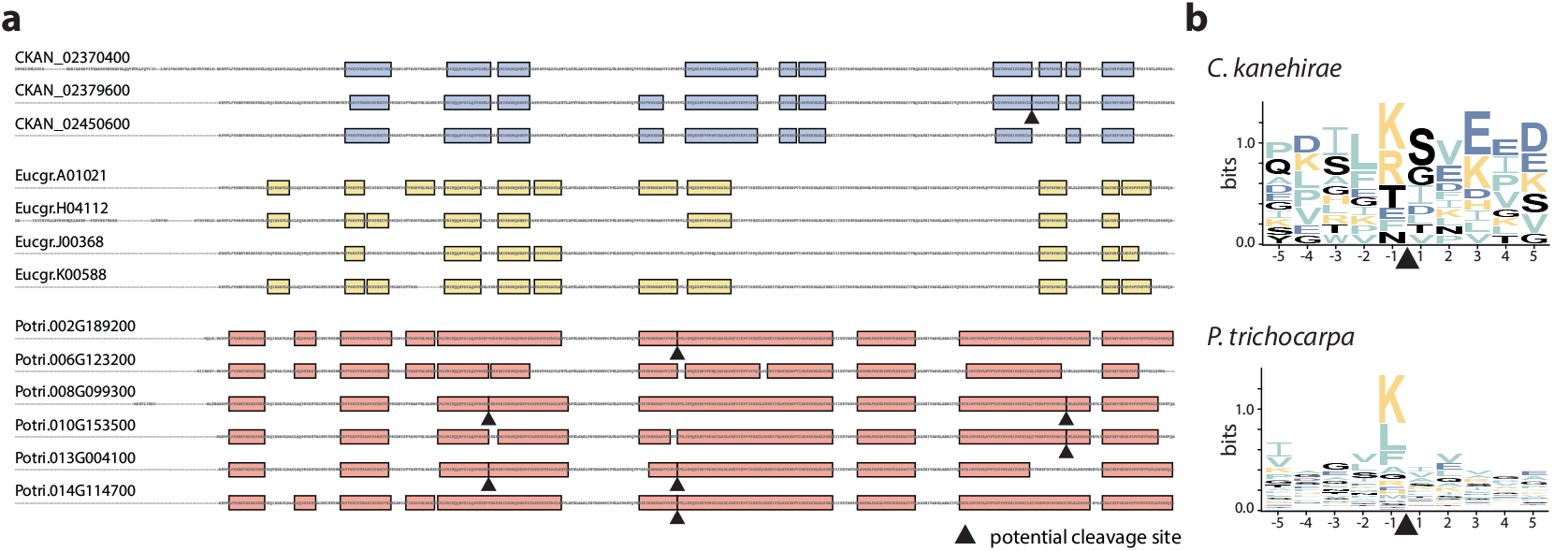
Conservation of sequence of sap peptide precursor genes. *(a)* Schematic of peptide-containing regions across species. The alignment illustrates protein sequences of an orthogroup shared among the three species, with colors indicating the regions containing peptides based on each species’ peptidome data. The expression levels of the corresponding genes are shown on the right. *(b)* Amino acid frequency at putative peptide cleavage sites. The motifs show the relative frequency of amino acids surrounding cleavage sites, with 89 unique sequences identified in *P. trichocarpa* and 12 in *C. kanehirae*, derived from 262 and 18 possible cleavage sites, respectively. Due to the limited peptide data, *E. grandis* was excluded from this analysis.

## Discussion

For over two centuries, the role of living xylem cells has largely been viewed as limited to preparing for their eventual function as dead, hollow structures responsible for fluid transport and mechanical support. This study challenges that long-standing paradigm by significantly expanding our understanding of the diversity of sap peptide precursor gene families. Whereas previous research identified approximately 50 representative families across species^14^, we report thousands of precursor genes in *Populus trichocarpa* and two other woody species. The total number of characterized peptides reached 7,039, marking a substantial expansion over the previously reported 1,997 sap peptides^34^. This substantial increase in peptide identification has allowed for a more comprehensive mapping of precursor genes, and, through transcriptome sequencing, we revealed that living xylem cells predominantly express these sap peptide precursors. This discovery positions living xylem cells as active and dynamic command centers for coordinating long-distance communication across the plant, and pointing to an evolutionarily conserved role in plant adaptation.

Traditionally, genes encoding small secreted peptides (SSPs) have been defined as proteins under 200–250 amino acids, often characterized by cysteine-rich motifs and post-translational modifications (PTMs)^14^. In the previously reported sap proteome in herbaceous plants^41^, about 70-80% of the sap proteins contained an N-terminal signal peptide. However, our analysis using LC-MS/MS-based peptidomic and transcriptomic evidence has enabled a direct experimental connection between sap peptides and their precursor genes. Many of these peptide precursor genes are non-Cys-rich, lack PTMs and conventional N-terminal signal sequences, tend to be more intrinsically disordered, and exceed the typical size threshold of 250 amino acids. These findings aligned with recent perspectives^14^ that suggest canonical SSP features may miss functionally significant and tissue-specific peptides, particularly those involved in long-distance signaling. Differences between *P. trichocarpa* and other species like *A. thaliana* and *M. truncatula* could arise from tissue specificity, environmental conditions, or functional roles, as some predicted SSPs might function as short-distance signals. Additionally, known families like CLE and CEP were only lowly expressed, indicating their roles may be limited to localized signaling. Comparisons with the *Glycine max* xylem sap peptidome further show context-specific variations, suggesting species-specific adaptations. These findings highlight that expanding SSP criteria is essential to capture the diversity and complexity of xylem-derived signaling peptides.

By integrating both vascular sap peptidome data and living xylem transcriptomes, our study provides new insights into the origin of long-distance mobile peptide precursors and their regulation. The high expression levels of these precursor genes in living xylem cells suggest that these cells are not merely passive but actively produce peptides crucial for signaling throughout the plant. This shifts the understanding of living xylem cells, previously regarded as serving only in secondary cell wall deposition and mechanical support^36,42^, towards their role in coordinating systemic communication. In support of this concept, the ‘good neighbor hypothesis’ proposes that living non-lignifying xylem cells provide numerous substrates necessary for neighboring lignifying cells, emphasizing a non-cell-autonomous process for cell wall formation^36,43^. Recent findings in Norway spruce demonstrate that monolignols, building blocks for lignin biosynthesis, are delivered to the cell wall via extracellular vesicles along with the enzymes for polymerization^44^. We propose that this cooperative behavior may also encompass the production of signaling molecules, with living xylem cells playing a dual role, contributing to an extensive network of intercellular communication. The ancestral functions of these precursor peptide genes are likely to be maintained by pervasive purifying selection. In addition, we further propose that these precursors may have also originated from other sources such as the degradome^33,39^, or by utilization short peptide sequences into housekeeping genes. Such evolutionary origins indicate that these peptides may have initially served as by-products of general cellular processes before being co-opted for specialized long-distance signaling functions. Morphological differences between species, including the diversity of xylem cell types, also suggest species-specific adaptations that shape the expression and function of these precursor genes. Taken together, these findings shed light on the evolutionary and functional origins of sap peptide precursors and highlight their essential role in plant communication and adaptation.

The identification of 331 and 899 sap peptides in *Eucalyptus grandis* and *Cinnamomum kanehirae*, respectively, remains below the numbers achieved in *P. trichocarpa*, where earlier peptidomic studies^34^ have been complemented by recent advances in mass spectrometry and an optimized sample collection pipeline in a model organism with comprehensive annotations. These advances suggest that the number of identified sap peptides will continue to increase as techniques improve and different conditions are examined^6,45^. Nevertheless, a limitation of our study was the integration of available datasets across species, highlighting the inherent challenges of working with non-model organisms. These discrepancies may be due to differences in experimental quality or intrinsic biological variation, such as morphological differences in wood formation. For example, in *P. trichocarpa*, the transition from primary to secondary growth occurs over a relatively short stem length, whereas in *E. grandis*, this process is more gradual and extended (**Supplementary Figure 7**). Such morphological differences are likely to contribute to the observed variability in expression profiles between sap peptide precursor orthologs and highlight the need for more refined experimental techniques to ensure consistency across diverse woody species.

In summary, this study provides the first comprehensive characterization of vascular sap peptides in multiple woody species, highlighting the previously underestimated roles of living xylem cells in both peptide production and long-distance plant-wide signaling. Our findings suggest that living xylem cells, traditionally viewed as structural and transport conduits, are in fact factories for long-distance peptides responsible for plant-wide signaling. The variation and evolutionary history of sap peptide precursors between species offer new insights into the complex regulatory networks governing plant development and stress responses. This expanded view of xylem function invites further elucidation of the functional roles of these peptides^34^ in plant adaptation, with promising applications for enhancing resilience to environmental stressors.

## Materials and Methods

### Vascular sap collection and peptide extraction

Ten-to twelve-month-old *Populus trichocarpa* plants with stem diameters of 1 to 1.5 cm, were used for vascular sap collection. The stems were cut approximately 2 cm above the soil, and the sap accumulating at the top of each stump was collected by pipetting over a period of 30-minutes. The stems were then segmented and centrifuged at 3,000 × g at room temperature to extract additional sap.

Peptide isolated from the sap sample was carried out using a solid phase extraction (SPE) column (Sep-Pak tC18, 1 g, 6 cc; Waters, part no. WAT036795). The SPE column was initially activated by 10 mL of 100% methanol and then equilibrated with 15 mL of 0.1% trifluoroacetic acid (TFA). The sap sample was then loaded onto the columns, followed by a washing step with 10 mL of 0.1% TFA. Peptides retained on the columns were eluted sequentially with 15 mL of 60% methanol and 15 mL of 80% methanol. The eluates from each methanol concentration were dried using a vacuum concentrator (miVac Duo Concentrator; Genevac) and then reconstituted in 100–400 μL of 0.1% TFA. The solution was centrifuged at 13,000 × g for 10 minutes at 4°C to remove any undissolved pellet, and the supernatant was collected. Desalting was performed using ZIPTIP® C18 pipette tips (Millipore), and peptides were eluted with 50% acetonitrile (ACN) in 0.1% TFA. The desalted samples were then dried again, reconstituted in 0.1% formic acid (FA), and centrifuged at 13,000 × g for 10 minutes at 4°C. The final supernatant was transferred to a sample vial for LC-MS/MS analysis.

### Sap peptidomic analysis using LC-MS/MS

The 60% and 80% methanol eluted sap samples were analyzed using an Easy nLC 1200 system coupled to an Orbitrap Fusion Lumos mass spectrometer (Thermo Fisher Scientific). Peptide separation was performed on an Acclaim PepMap 100 C18 trap column (75 µm × 2.0 cm, 3 µm, 100 Å; Thermo Fisher Scientific) followed by an Acclaim PepMap RSLC C18 nano LC column (75 µm × 25 cm, 2 µm, 100 Å). The mobile phase consisted of 0.1% formic acid (FA) in ultra-pure water, with an elution buffer of 0.1% FA in 100% acetonitrile (ACN). Peptides were eluted over a 90-minute linear gradient of 5% to 25% ACN/0.1% FA at a flow rate of 300 nL/min. In data dependent acquisition (DDA) mode, a full scan was conducted from m/z 350–1600 at a resolution of 120,000 (at m/z 200), followed by MS/MS fragmentation within a 3-second cycle. The maximum ion injection time for the full scan was set to 50 ms. MS/MS acquisition was carried out with a 1.4 Da isolation window, an AGC target of 50,000, 35% normalized collision energy, a maximum injection time of 120 ms, and a resolution of 15,000.

### Identification and functional annotation of sap peptide precursor genes

For *P. trichocarpa* peptidomic profiling, we generated a target-decoy database for MS/MS ion searching and evaluated the false discovery rate (FDR)^46^, for identifying confident peptides. The database generated by combining the protein sequences of *P. trichocarpa* with the randomized protein sequences (decoy) using Trans Proteomics Pipeline (TPP) version 5.1^47^. The protein sequences of *P. trichocarpa* (*Populus trichocarpa* v4.1) were downloaded from Phytozome^48^. The MS raw data of peptidomic dataset obtained from LC-MS/MS was first converted into mzXML format using MSConvert^49,50^, and then processed by UniQua with optimized parameters (Smoothing Width: 5, Centroiding, Deisotoping, Noise Removal, Intensity Cutoff: 500)^51^. The UniQua processed data were converted into Mascot generic format (.mgf) by MSConvert^50^. The processed mgf files were searched against a concatenated target-decoy database for corresponding species (described above) using Mascot search engine (version 2.3; Matrix Science, London, UK) without specifying enzyme cleavage rules. The mass tolerances used in the MS/MS ion search for peptide precursors and fragments were 10 ppm and 0.05 Da, respectively. The oxidation of methionine and the sulfation of tyrosine were considered as the variable modifications. Three independent scoring methods, Mascot, Delta Score (DS), and Contribution Score (CS)^32^ were used to evaluate the peptide identification based on FDR. The Mascot search result files (.dat) were used to extract the Mascot scores and to re-calculate the DS and CS based on Mascot scores using the CScore program^32^. The identified peptides with FDR below 0.05 from at least one of the three scoring methods were regarded as positive hits.

Vascular sap peptides were mapped to the proteome. Among the 7,039 identified peptides, 7,021 were mapped to the longest protein isoform and selected for subsequent analysis. Overlapping regions were consolidated into single contiguous segments and defined as peptide-containing regions for subsequent analysis. Protein family domains of the precursor genes were identifed using pfam_scan (ver. 1.8)^52^, querying the Pfam database (ver. 37)^41^. The N-terminal signal sequence within each protein was predicted using SignalP (v6.0)^53^. Gene ontology terms were annotated using eggNOG-mapper (v2)^54,55^. In addition, disordered protein regions and disordered binding regions were predicted using IUPred2A and ANCHOR2^56,57^, respectively.

### Orthogroup inference and comparative genomics

Protein sequences of 14 plant species were downloaded from Phytozome^58^ (*Zea mays* Zm-B73-REFERENCE-NAM-5.0.55, *Vitis vinifera* v2.1, *Solanum lycopersicum* ITAG5.0, *Populus trichocarpa* v4.1, *Oryza sativa* v7.0, *Musa acuminate* v1, *Mimulus guttatus* v2.0, *Glycine max* Wm82.a2.v1, *Eucalyptus grandis v2*.*0, Daucus carota* v2.0, *Cinnamomum kanehirae*^59^ v3, *Aquilegia coerulea* v3.1, *Arabidopsis thaliana* Araport11, *Amborella trichopoda* var. SantaCruz_75 HAP2 v2.1). Gene families or orthogroups (OGs) were inferred from the longest protein isoforms using Orthofinder (v2.5.5)^60^. For orthogroups containing genes from at least three species, protein sequences were aligned using MAFFT (v7.525)^61^, and a phylogenetic tree was constructed using VeryFastTree (v4.0.2)^62^. A species tree was inferred from 10,808 OG gene trees using ASTRAL-Pro (v1.10.1.3)^63^.

Datasets of small secreted proteins (SSPs) and peptide cleavage products (PCPs) from *A. thaliana, G. max, P. trichocarpa*, and *M. truncatula* were obtained from PlantSSP^16^, REF10^10^, REF33^33^, and MtSSPdb^17^, respectively. As these datasets used gene identifiers from older genome references, we updated the sequence identifiers by aligning the corresponding protein sequences to the latest genome assemblies using DIAMOND (version 2.1.9) ^64^. We then assessed the overlap between these datasets and our peptidomic data by determining whether the orthogroups contained members from both the available datasets and the putative *P. trichocarpa* sap peptide precursor genes.

### Sampling for laser capture microdissection

The *P. trichocarpa* plants were grown in a walk-in growth chamber under 16-hour light/8-hour dark cycles at temperatures between 24°C and 26°C. For *E. grandis*, trees were planted in a walk-in greenhouse under sunlight and temperature of 20–30°C. Tree stems were collected from plants approximately 180 cm tall. The 3rd, 4th, 5th and 8th internodes of the *P. trichocarpa* tree stems were used. The 4th internodes of *P. trichocarpa* were separated into two parts, T1 and T2, which were located next to the 3rd and 5th internodes, respectively. Each biological replicate in *P. trichocarpa* was under a similar growth rate such that the xylem bundles in T1 were not connected, while the xylem of T2 was connected to form a circular pattern **(Figure 2c)**. The 3rd, 5th and 8th internodes were defined as P, S1 and S2, respectively. For *E. grandis*, the 13th, 14th, 15th and 18th internodes were harvested. The 14th and 15th internodes of each biological replicate in *E. grandis* were under the condition that the xylem bundles in the part of the 14th internode adjacent to the 13th internode (designated as T1) remained separated, while the xylem in 15th internode was organized into a circular structure (**Supplementary Figure 7**). The part of the 14th internode adjacent to the 15th internode was defined as T2, and the 13th, 15th and 18th internodes were labeled as P, S1 and S2, respectively.

The 1 mm thick segments sampled from the 3rd, 4th, 5th and 8th internodes of *P. trichocarpa* and the 13th, 14th, 15th and 18th internodes of *E. grandis* were fixed in 100% acetone (Sigma-Aldrich, USA) and vacuumed for at 4°C for 45 minutes. The segments in 100% acetone were then incubated at 4°C overnight and at 37°C for 1.5 h. The segments were placed in a graded n-butanol (ECHO chemical, Taiwan) series (30, 50, 70, 90, and 100%, v/v) diluted with 100% acetone at 58°C for 20 minutes in each solution followed by a graded paraffin (Leica, USA) series (50, 60, 80, and 100%, v/v) diluted with 100% n-butanol at 58°C for 30 minutes in each solution. The 16-μm sections were prepared from the paraffin embedded segments using a rotary microtome (Microm, HM355E). The embedded paraffin in the sections was removed with xylene (Sigma-Aldrich, USA) for 30 minutes at RT, followed by 99.5% ethanol (ECHO chemical, Taiwan) for 30 minutes at RT.

### Laser capture microdissection

Segments cut from the 3rd, 4th, 5th and 8th internodes of *P. trichocarpa* and the 13th, 14th, 15th and 18th internodes of *E. grandis* were immediately frozen in liquid nitrogen and stored at −80°C. The segments were fixed on cryostat chucks with Tissue-Tek O.C.T. Compound (Sakura Finetek, USA) and used for sectioning with 16-μm thickness using a cryostat (Leica CM1900) at −20°C. The PET membrane of metal frame slides (Leica, USA) was covered with 99.5% ethanol (ECHO chemical, Taiwan) and then the cryosections were placed on the membrane for dehydration. Four biological replicates of *P. trichocarpa* and *E. grandis* for each developmental stage were collected from an area of approximately 1,500,000 μm^2^ of cryosections within approximately 20 cell layers using laser microdissection systems (Leica LMD7000). Dissected cells were collected within 30 minutes in 500-μL eppendorf caps containing a mixture of 50 μL RLT buffer (QIAGEN, USA) and 1% beta-mercaptoethanol (Sigma-Aldrich, USA). Laser microdissected samples were snap frozen in liquid nitrogen and stored at −80°C.

### RNA sequencing

Total RNA from dissected cells was extracted using RNeasy Plant Mini Kit (QIAGEN, USA) according to the user manual. The mRNA of each developmental stage was amplified using the Arcturus® RiboAmp® HS PLUS RNA Amplification Kit (Applied Biosystems, USA) according to the manufacturer’s instruction. The sequencing library of each sample was constructed from the amplified mRNA using the NEBNext® Ultra™ II Directional RNA Library Prep Kit (New England Biolabs, USA) according to the user guide. The RNA-seq libraries of *P. trichocarpa* and *E. grandis* were sequenced on HiSeq X Ten platform (BGI Genomics Co., Ltd, China and Novogene Co., Ltd, China, respectively) to obtain 150-bp length of paired-end reads.

### Microscopic imaging

Stem sections were visualized under an Olympus BX53 upright microscope in bright field and with an excitation filter range of 360–370 nm and an emission filter range of 420–460 nm to observe lignin fluorescence. Images were acquired using the Olympus cellSens platform. Sections for LCM were observed using the camera function integrated in the laser microdissection systems (Leica LMD7000).

### Tissue transcriptome datasets

The tissue transcriptomes of *Populus trichocarpa, Eucalyptus grandis* and *Arabidopsis thaliana* were downloaded from NCBI under accession numbers PRJNA320431, PRJNA731920 and PRJNA595603, respectively.

The 12-month-old stems of *C. kanehirae*, grown from a 10-year-old tree stump, were used for tissue transcriptome analysis. Four tissues, leaves, young shoots, phloem and xylem, were collected. Young shoots were harvested as the first to third internodes for RNA extraction and the first to eighth internodes for protein extraction. Bark peeled from the stem served as the phloem sample, while stem-differentiating xylem was obtained by scraping the surface of debarked stems. Four tissues (xylem, phloem, leaves, and young shoots) were used for RNA extraction by the CTAB method, utilizing a modified CTAB extraction buffer based on REF65^65^. The buffer contained 2% (w/v) hexadecyltrimethylammonium bromide, 0.1 M Tris-HCl (pH 9.0), 25 mM EDTA (pH 8.0), 2 M NaCl, and 1% (w/v) PVP-40, with 2% (v/v) β-mercaptoethanol and 50 mM ascorbic acid were added just before use. Each tissue sample was ground to a fine powder in liquid nitrogen, and 0.5 g of each powdered sample was transferred to 15 mL tubes. Each sample was resuspended in 5 mL of pre-warmed (65°C) CTAB extraction buffer, vortexed thoroughly, and incubated at 65°C for 10 minutes. After centrifugation at 12,000 × g for 5 minutes at room temperature, the supernatant was transferred to fresh 15 mL tubes, mixed with an equal volume of chloroform alcohol (24:1), vortexed, and centrifuged at 12,000 × g for 10 minutes at room temperature. This chloroform extraction step was repeated once. The aqueous phase was combined with one third volume of 8 M lithium chloride and stored overnight at 4°C. RNA pellets were then collected by centrifuging at 12,000 × g for 20 minutes at 4°C and further purified using the Qiagen RNeasy Plant RNA Isolation Kit. RNA libraries were constructed using the NEBNext® Ultra™ II Directional RNA Library Prep Kit for Illumina®, followed by RNA sequencing on the Illumina Novaseq 6000 platform.

### Transcriptome analysis

Raw RNA-Seq reads were trimmed using fastp (v0.23.2)^66^ to remove adaptor sequences and low-quality sequences. The cleaned reads were then mapped to the respective genome using STAR (v 2.7.10b)^67^ and gene counts were generated with featureCounts (v 2.0.3) ^68^. Gene expression levels were quantified as transcripts per million (TPM) for each sample, and the median TPM values across replicates were used as representative gene expression measures for cross-species comparisons. Gene ontology (GO) enrichment analysis was conducted using TopGO (v2.56.0)^69^. For each orthogroup, TPM values of all member genes within a species were summed per sample and converted into proportions. Median proportions across replicates were used to represent the transcriptome contribution of each orthogroup.

To assess the conservation trend of gene expression levels across species, we categorized the expression levels of orthogroups into quantiles within each species. Specifically, we divided the orthogroups into quantiles based on their expression levels, assigning them to categories such as high, medium, and low expression. We then evaluated changes in these expression categories between species by comparing the assigned quantiles in pairwise species comparisons (e.g., *Eucalyptus grandis* vs. *Populus trichocarpa*, and *E. grandis* vs. *Cinnamomum kanehirae*). For each comparison, we calculated the frequency with which orthogroups maintained their expression category across species. In the *E. grandis*–*P. trichocarpa* comparison, highly expressed orthogroups remained in the same expression category 2.30 times more frequently than other orthogroups. In the *E. grandis*–*C. kanehirae* comparison, this factor was 1.98. When focusing specifically on orthogroups containing sap peptide precursor genes, these differences were even more pronounced—3.93 times in the *E. grandis*–*P. trichocarpa* comparison and 3.22 times in the *E. grandis*–*C. kanehirae* comparison. These analyses revealed fewer shifts in the highest expression categories between species, indicating that expression changes are more constrained for highly expressed orthogroups, particularly those encoding sap peptide precursor genes.

### Analysis of sequence conservation

For each orthogroup, the most highly expressed ortholog from each species was selected. Protein sequences from *P. trichocarpa, E. grandis*, and *C. kanehirae* were aligned using MAFFT (v7.525)^61^. Codon alignments were obtained from inputting nucleotide sequences and protein sequence alignments using PAL2NAL (v14)^70^. The ratio of nonsynonymous to synonymous substitution rates (*d*N/*d*S ratio) between *Populus trichocarpa* and two other species was estimated using the Codeml program in PAML (v4.10.7)^71^. trimAl (v1.5.rev0)^72^ was used to remove excessive gaps, minimizing the underestimation and ensuring a more accurate assessment of the identity of the protein alignments. Sequence identity was calculated ignoring gaps that were present in both species compared.

### Identification of putative cleavage sites

Adjacent, non-overlapping peptide-containing regions were designated as potential peptide cleavage sites, defined by considering five amino acid residues both upstream and downstream of the boundary site. To minimize gene copy bias, only the most highly expressed member of each orthogroup was selected. Selected sequences were deduplicated using usearch (v11.0.667)^73^ and visualised using WebLogo (v3.0)^74^.

## Data Availability

All sequences generated from this study were deposited on NCBI under BioProject PRJNA1180722. Accession numbers of individual samples can be found in **Supplementary Table 9**. All peptidomic datasets were deposited to the ProteomeXchange Consortium via the PRIDE partner repository with the identifier PXD032088^34^ and PXD057644.

## Acknowledgement

We thank Yu-Ching Liu for uploading the RNAseq data to NCBI and Dr. Shu-Ming Tsao (ez-Omics Co., Ltd.) for target-decoy FDR calculation. The mass spectrometry analyses were supported by Dr. Chuan-Chih Hsu (Proteomics Core Laboratory, Institute of Plant and Microbial Biology, Academia Sinica).

## Author contributions

IJT, YCL and YLC conceived the study; PCL and JHY carried out the sap collection. YLC and PCL carried out the sap peptide extraction. PCL carried out tissue RNA extraction. CHC and YLC carried out the peptidomic identification.; YCL, CCT, CEH and CYL carried out the xylem laser capture microdissection and transcriptome sequencing; IJT and MRL carried out the comparative genomics and transcriptomics. MRL, YCL, YLC and IJT wrote the manuscript with input from all authors. All authors read and approved the final manuscript.

