## Supplementary Info for "Living xylem cells encode a large number of conserved gene families responsible for vascular sap peptides"

**Supplementary Figure 1. Length distribution of sap peptides and proteins in *Populus trichocarpa*. (a)** Length distribution of 7,021 peptides that were mapped to longest isoform of *P. trichocarpa* proteome. **(b)** Length distribution of sap peptide precursor proteins compared to the entire *P. trichocarpa* proteome. The dashed lines represent the median length of each distribution.

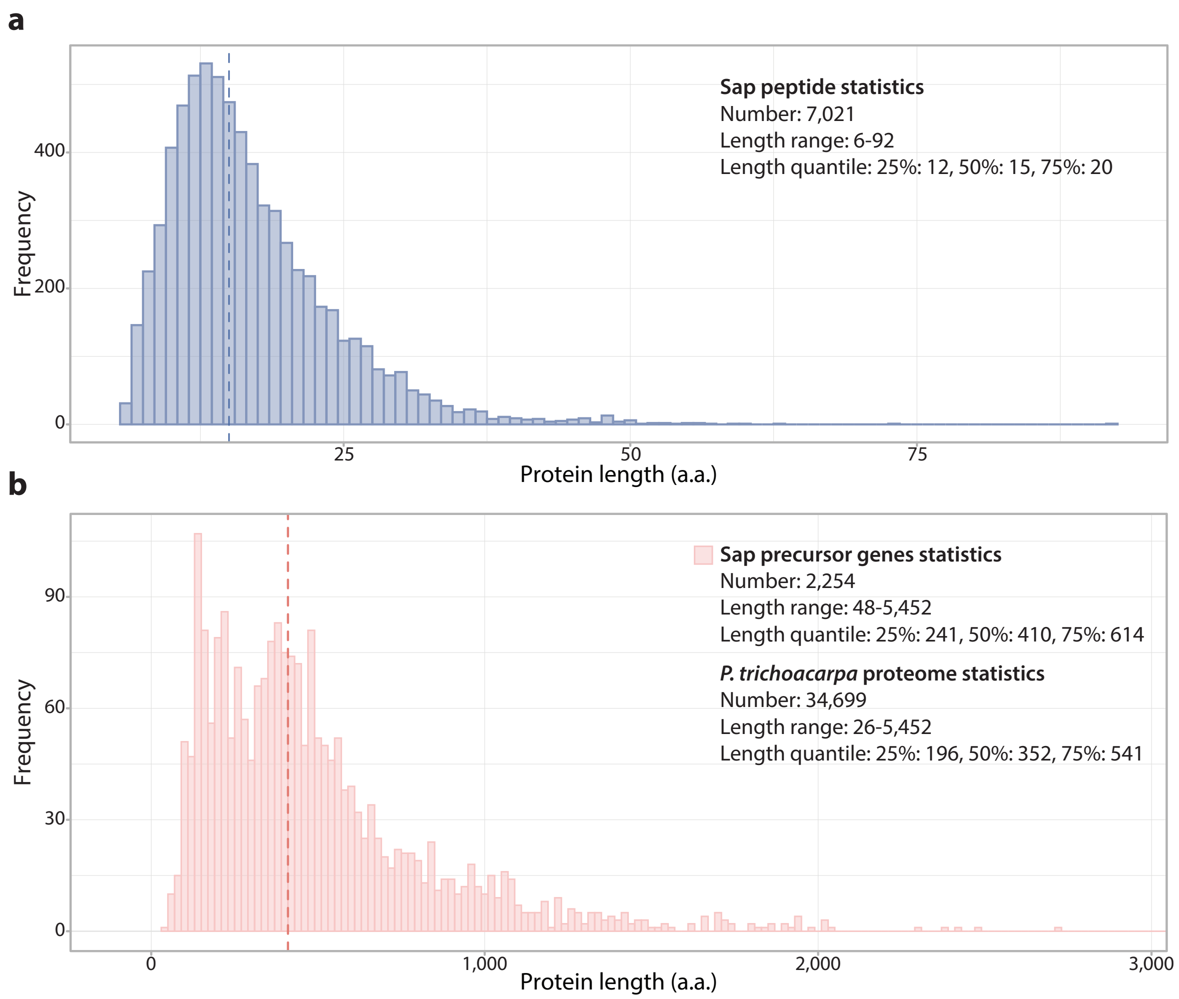

**Supplementary Figure 2. Distribution of the number of orthogroups of randomly selected genes.** The histogram represents the distribution of the number of orthogroups observed by randomly selecting 2,254 genes from the entire gene set in *P. trichocarpa*, with 10,000 iterations. A singleton was counted as one orthogroup. The brown dashed line represents the median of the distribution, while the black line denotes the observed number of orthogroups for the sap precursor genes. P value was inferred from obtaining the observed value from the distribution.

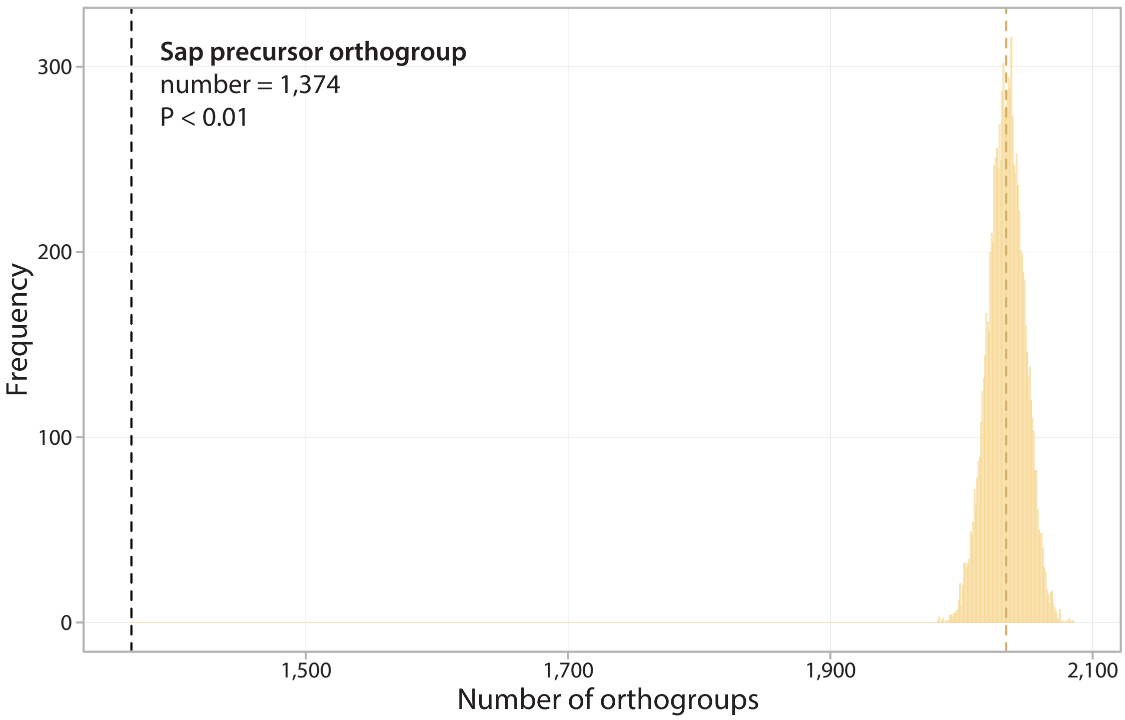

**Supplementary Figure 3. Characteristics of the sap peptide precursor genes.** (a.) Distribution of the total number of genes and the number of genes containing sap peptides within 1,335 orthogroups and 39 singletons in *P. trichocarpa*. Colors represent whether all genes in an orthogroup contain sap peptides (complete, in blue) or only some genes do (partial, in green). (b.) Proportion of orthogroups containing sap peptides in a random selection of 2,254 sap peptide precursor gene sets. The x-axis represents the number of randomly selected genes, ranging from 1 to 2,254, while the y-axis indicates the proportion of orthogroups in which selected genes contain sap peptides. Mean values from 100 bootstraps are plotted, with colors indicating orthogroup status—partial (green) or complete (blue)—to illustrate that all genes within an orthogroup are likely to be encode sap peptides as the number of detected peptides increases.

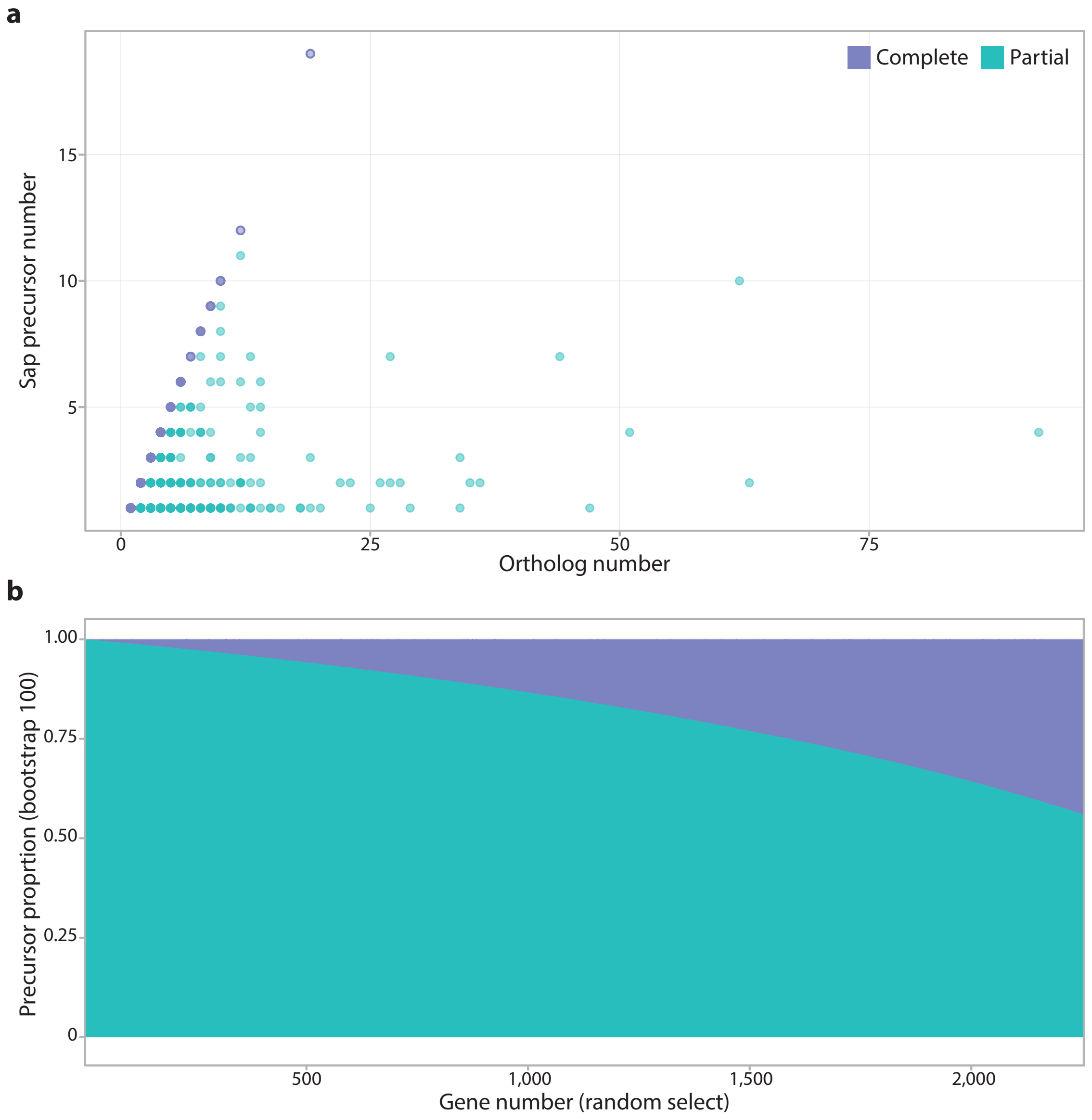

**Supplementary Figure 4. Intrinsic disorder of *P. trichocarpa* genes.** (a) Comparison of disordered regions and disordered binding regions across three gene categories: non-sap precursor genes, sap precursor genes with signal peptides, and sap precursor genes without signal peptides. Disorder scores were calculated for each amino acid using IUPred2A and ANCHOR2, with the proportion of amino acids scoring above 0.5 considered as disordered. Significant differences between categories are indicated (Wilcoxon ranked sum test; ****P < 0.0001; *P < 0.05, Wilcoxon test). (b) Spearman correlation analysis between the proportion of peptide-containing regions and protein disorder levels within *P. trichocarpa* genes. Sap precursor genes were categorized according to the presence (red) or absence (pink) of signal peptides.

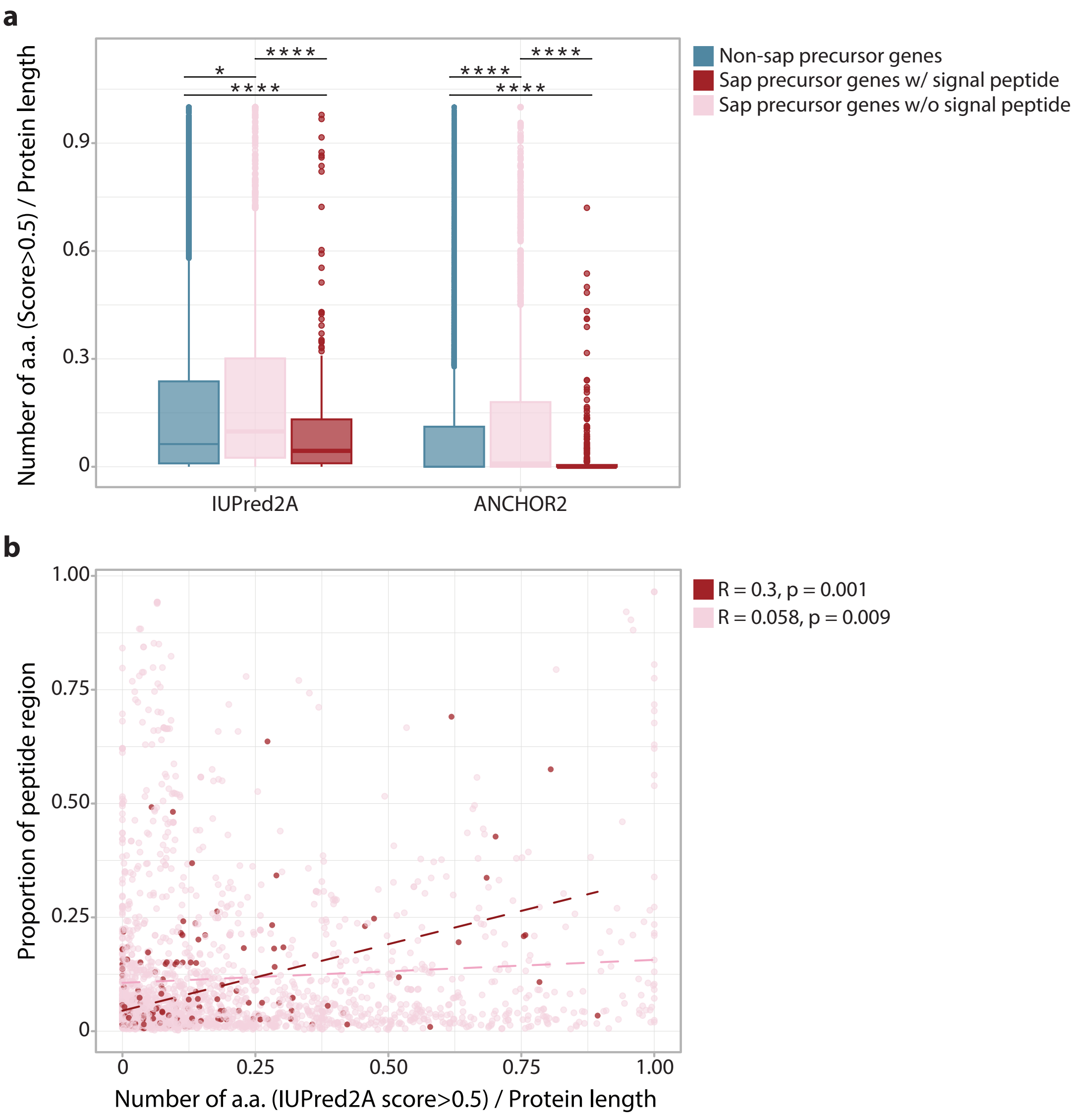

**Supplementary Figure 5. Comparison of inferred peptide precursor genes across studies.** A total of 4,804 sap precursor genes identified in our study were compared with predicted (mostly *in silico*) peptide precursor genes from different databases. Both overlapping orthogroups and genes within the orthogroups of each category are presented.

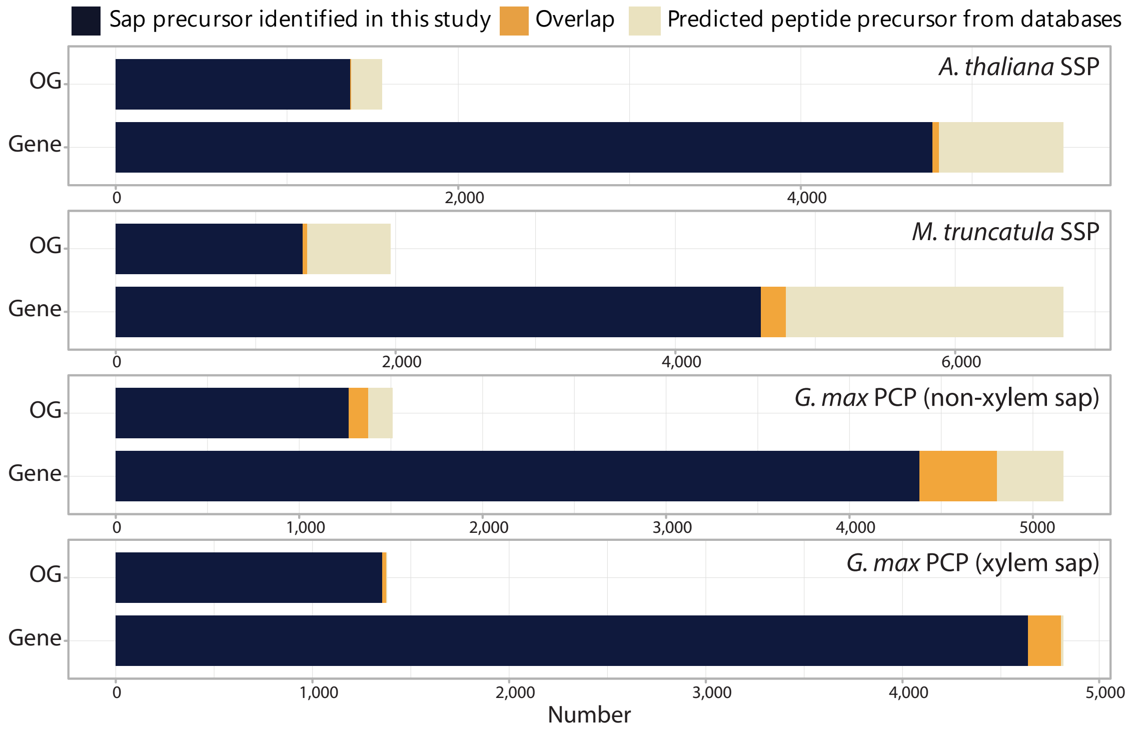

**Supplementary Figure 6. Examples of XAP family orthology between *G. max* and *P. trichocarpa*.** Sequence alignments were performed for (a) XAP1 and (b) XAP5A genes in *G. max* together with their corresponding orthologs in *P. trichocarpa*. Peptides of *G. max* and *P. trichocarpa* are highlighted in purple and yellow, respectively. Matched peptide residues are highlighted in red.

**a**

```
                                  65           75           85           95           105
Glyma19g29590.1|XAP1    LDHVVGNGEVVLVDMNEGFMERRVDLETQDYEGTGANKDHDPKSPGGA----

Potri.013G066600        QDAVISTDGEMLIDAGEGYIEGRMDLESTDYPGTGANNHHDPKTPGKA----
Potri.008G087700        QHQTITKR-----------GAGRENLEYTDYSGTGPNNRHTPEPPSGQGGN-
Potri.015G062300        THLVLQSKKK---DHDEHIAHGRKIVELNDYPGSGANNRHTPRPQFGRCVDC
```

**b**

```
                                  65           75           85           95           105          115          125
Glyma04g41240.2|XAP5A   LILIPL-SSGLAEGFGENMHPTHGLLYKDGIKM-NSRKLLVHDFVLDYDEAGPNPRHTKK------PGKGP

Potri.015G010500        LLLAPLSSSGLVEGFKEGMHP-HNSFVKDGIHMINARKLLL-D-MLDYGDAGANHKHDPRGKAVVVGGKSP
Potri.012G017400        LLLAPF-SSGFVEGFDEGMNS-YPSLHKDGIQV-NTRKLLLVD-ELDYDDAGANRRHDPRGRPGVGGYKNP
```

**Supplementary Figure 7. Anatomy during wood formation in *P. trichocarpa* and *E. grandis*.** The anatomical structures observed during different stages of wood formation are shown.

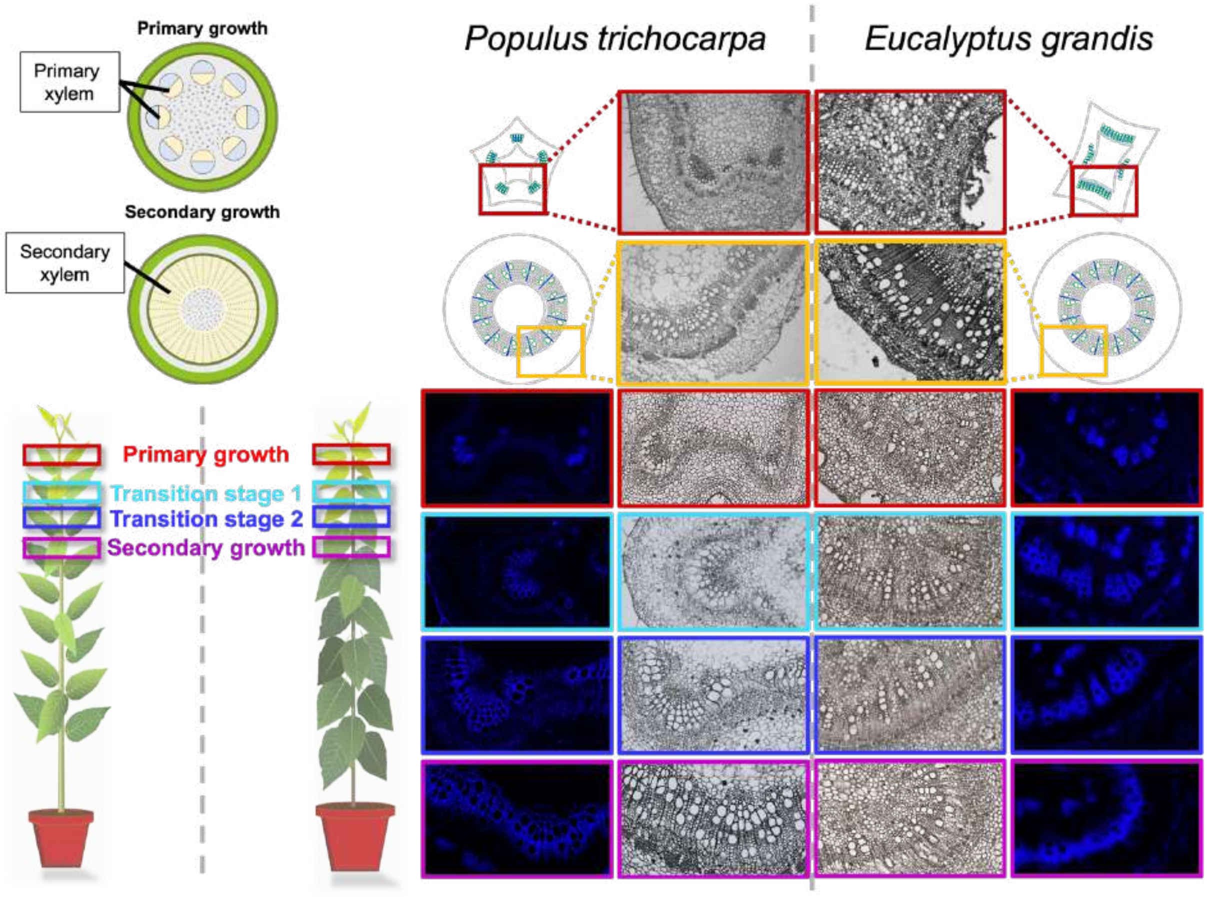

**Supplementary Figure 8. Cumulative expression of sap precursor genes across developmental stages in *P. trichocarpa*.** Cumulative expression profile of 4,804 sap precursor genes identified in *P. trichocarpa*. Colored lines denote different developmental stages, while dashed and dotted lines indicate the sum of TPMs from genes accounting for 50% and 75% of the total transcriptome, respectively, at each stage.

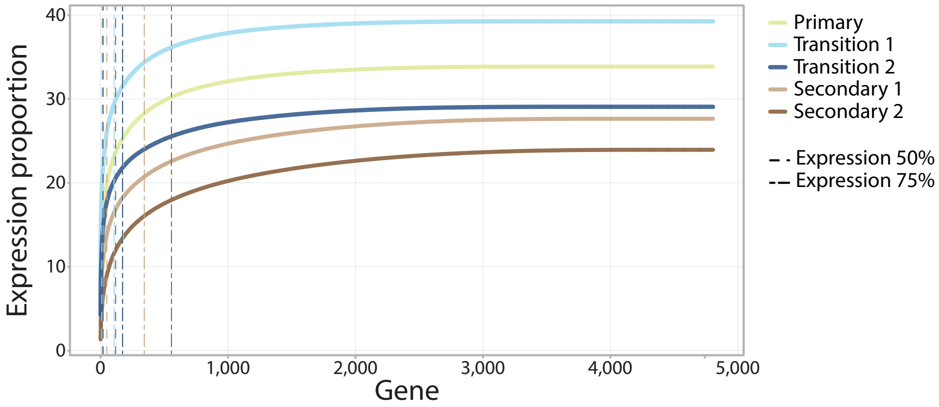

**Supplementary Figure 9. Expression profiles of XAP orthologs in *P. trichocarpa*.** Expression of *P. trichocarpa* orthologs within orthogroups of XAP (Xylem sap-Associated Peptides) genes from *G. max* are shown. Heatmaps show the expression levels (log2 TPM) of these XAP orthologs across different tissues and developmental stages. Stars indicate *P. trichocarpa* genes producing sap peptides.

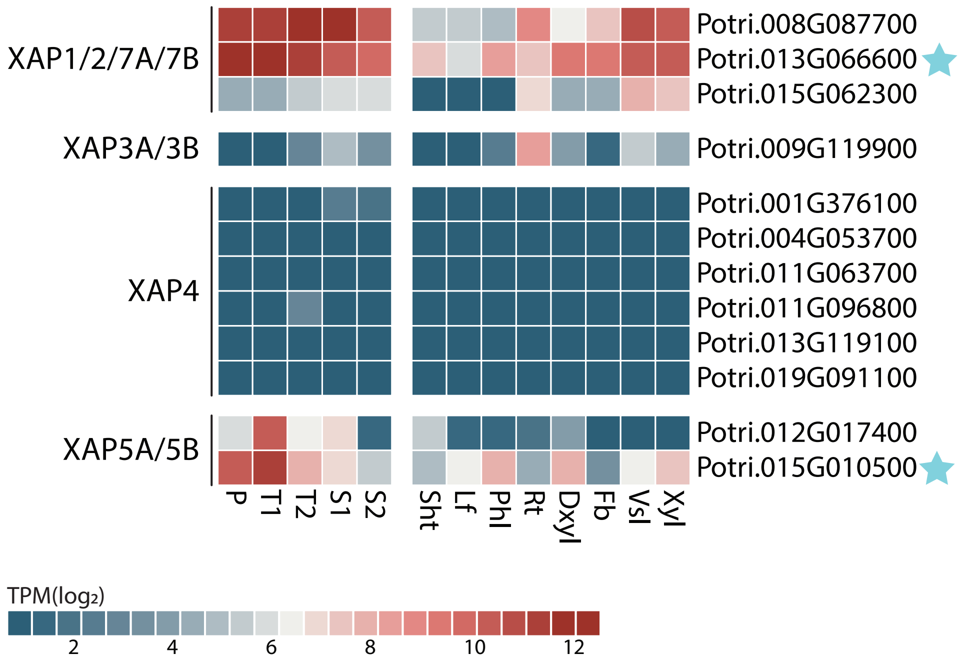

**Supplementary Figure 10. Relationship between protein length and non-overlapping peptide-containing region across xylem developmental stages in *P. trichocarpa*.** Each panel represents a different xylem development stage (primary, transition 1, transition 2, secondary 1, and secondary 2). The x-axis shows the length of the sap precursor proteins, and the y-axis shows the total length of the non-overlapping peptide-containing regions within each gene. Dot sizes indicate expression levels (TPM) and colors represent the number of different peptide regions within each precursor.

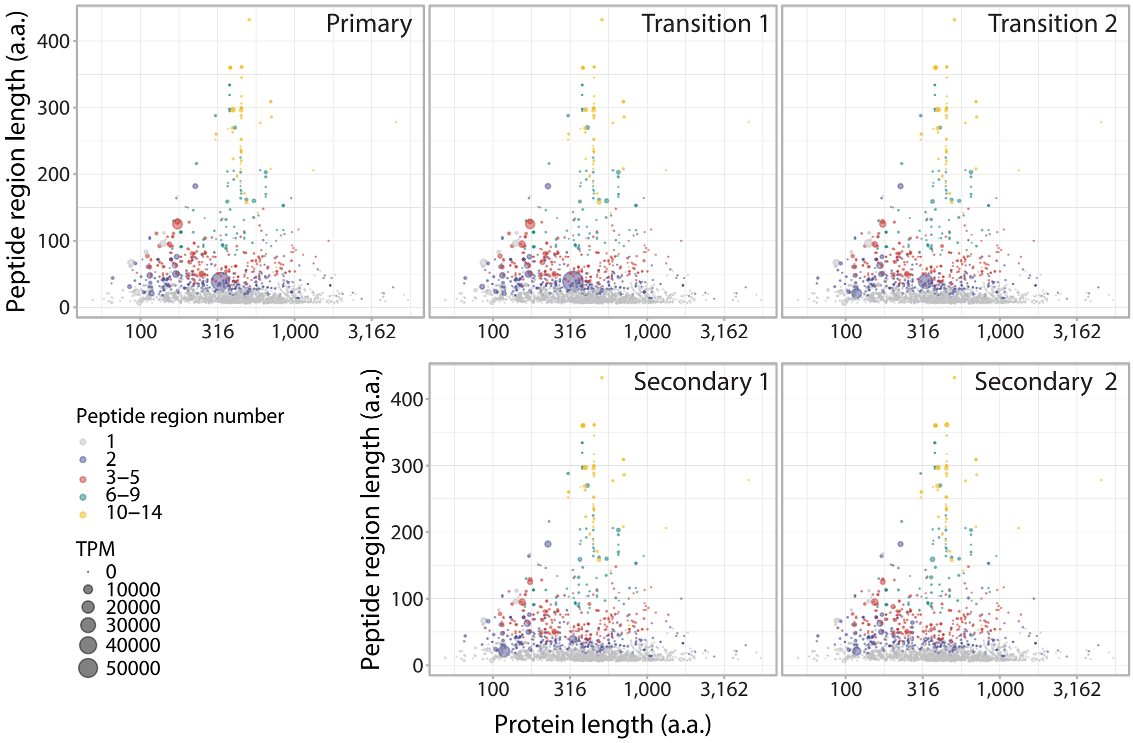

**Supplementary Figure 11. Expression patterns of sap peptide precursors across developmental stages and peptide characteristics.** Each panel categorizes precursor proteins by length and shows their contribution to the transcriptomes at different developmental stages (P, T1, T2, S1, S2). Colors represent the relative position of the peptide-containing regions within the precursor protein: N-terminal (red), balanced (green), and C-terminal (blue). Peptide regions that occupy more than 70% of themselves in either terminal are classified as N- or C-terminal, while those that are more evenly distributed are termed balanced.

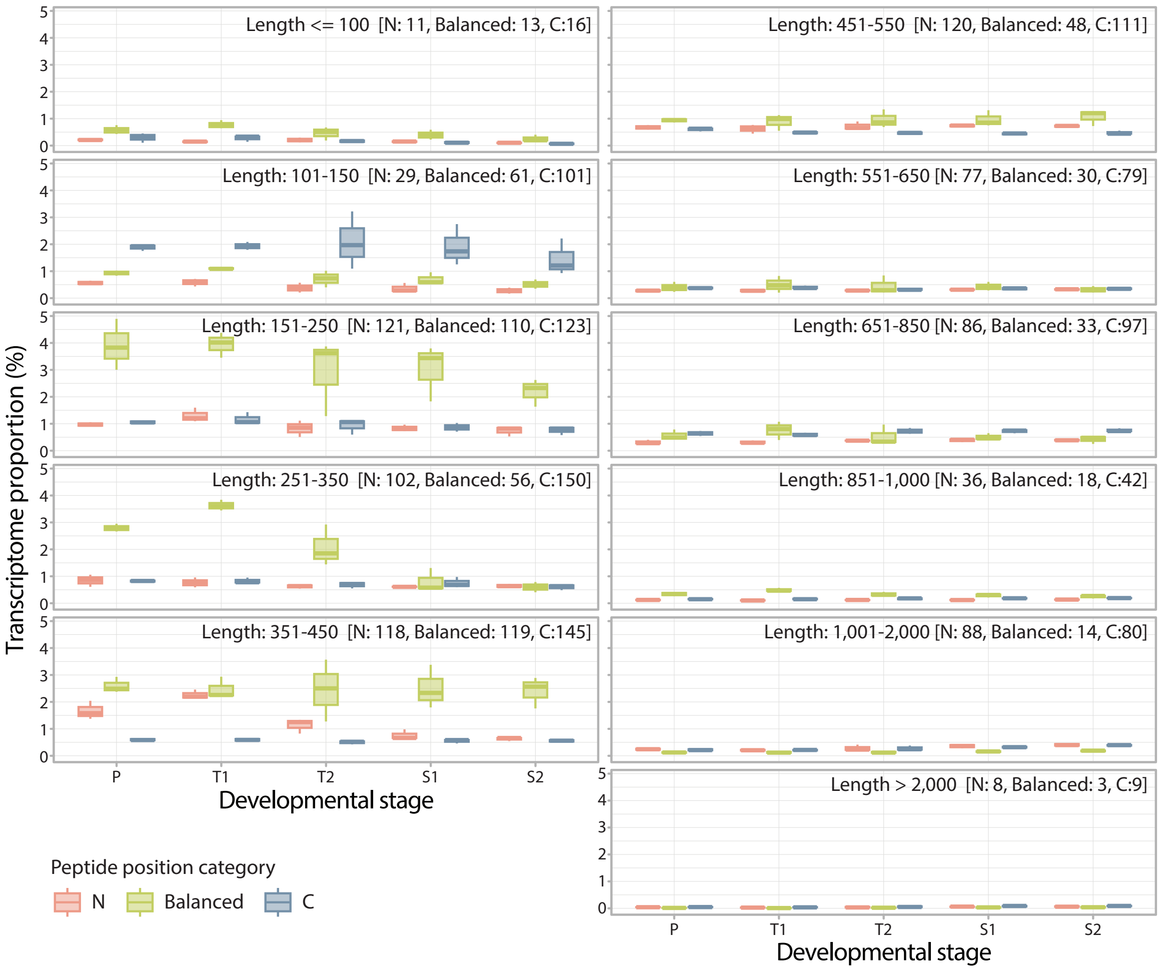

**Supplementary Figure 12. Cumulative expression of sap precursor genes across developmental stages in *E. grandis*.** (a) Expression profile of *E. grandis* precursor genes, based on orthologs identified from the *P. trichocarpa* peptidome. (b) Expression profile combining *P. trichocarpa* orthologs and precursor genes identified directly from the *E. grandis* peptidome. Colored lines denote different developmental stages, while dashed and dotted lines indicate the sum of TPMs from genes accounting for 50% and 75% of the total transcriptome, respectively, at each stage.

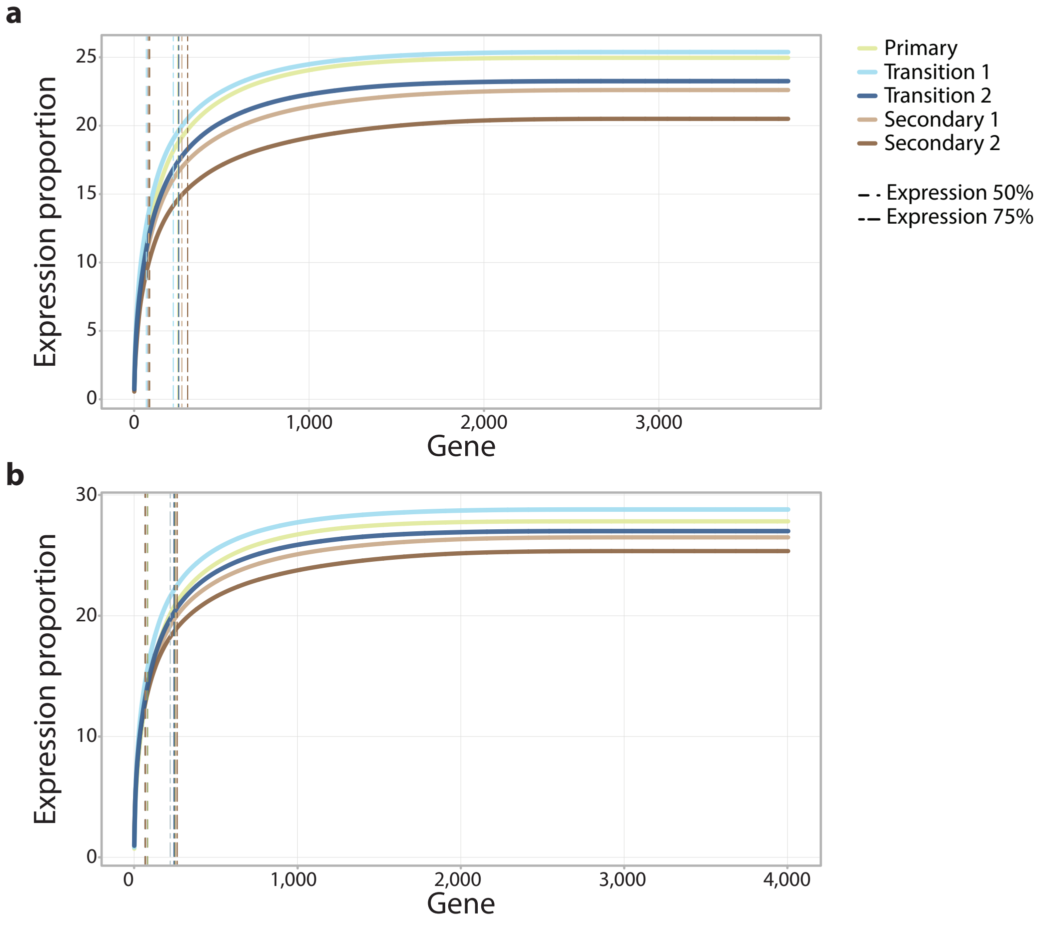

**Supplementary Figure 13. Expression patterns of sap peptide precursor genes across developmental stages in *E. grandis*.** (a) A total of 3,739 sap peptide precursor orthologs in *E. grandis* were inferred based on the *P. trichocarpa* peptidome. (b) 4,004 genes were identified by combining data of *P. trichocarpa* sap precursor orthologs and precursor genes from the *E. grandis* peptidome. Only genes with an expression value greater than 0.1% of the transcriptome are included. Genes are sorted by their expression levels in the T1 stage, and the top 10 expressed genes in each developmental stage are highlighted with different colors. Labels indicate the percentage transcriptome contribution for each highlighted gene across the stages (P, T1, T2, S1, S2).

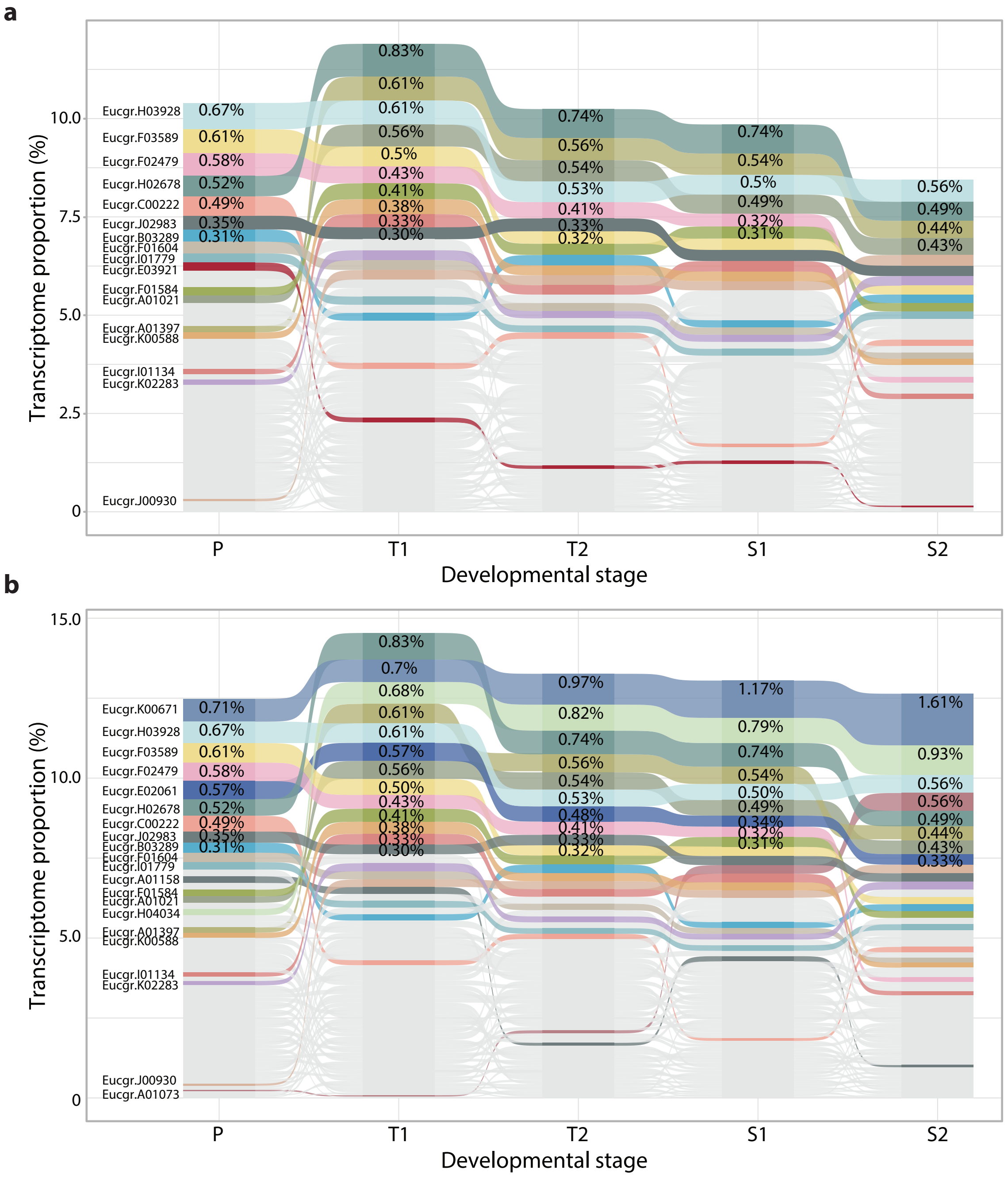

**Supplementary Figure 14. Correlation of gene expression between species.** (a) Comparison of gene expression (TPM in $log_2$ scale) between the T1 stage of *P. trichocarpa* and *E. grandis*, with correlation values calculated for all orthogroups (light blue) and specifically for sap precursor-containing orthogroups (dark blue). (b) Comparison between the T1 stage of *P. trichocarpa* and the xylem of *C. kanehirae*, showing correlation values for all orthogroups (light green) and sap precursor-containing orthogroups (dark green). Each point represents an orthogroup, with darker colors highlighting orthogroups containing sap precursor genes. Dashed lines indicate linear regression fits for each dataset.

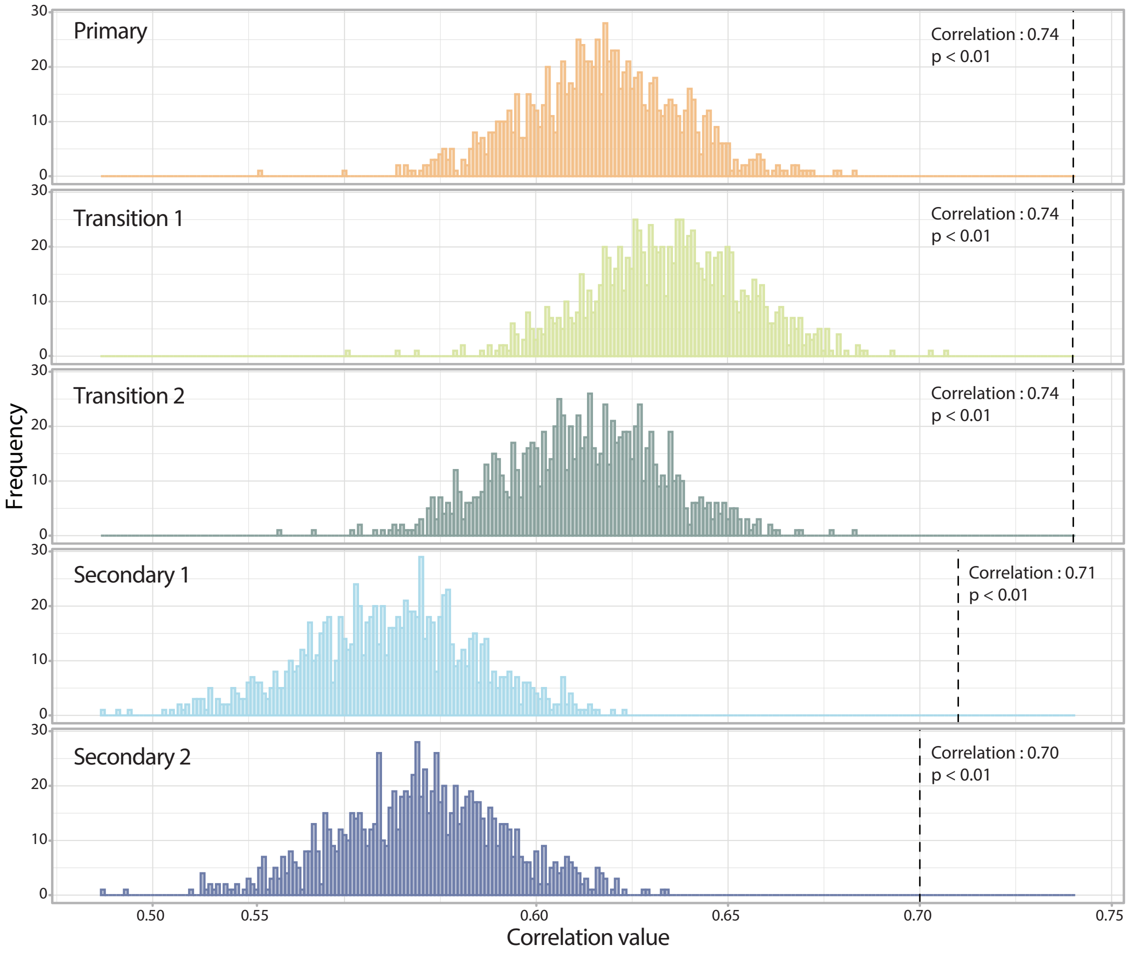

**Supplementary Figure 15. Distribution of the expression correlation values of between *P. trichocarpa* and *E. grandis* across developmental stages.** The distribution of Spearman correlation values was calculated between the expression of T1 stage in *E. grandis* across xylem developmental stages in *P. trichocarpa* from 1,223 randomly selected orthogroups over 1,000 iterations. The black dashed line marks the correlation of the expression of sap peptide precursors expression at T1 stage in *E. grandis* compared to their orthologs across xylem developmental stages in *P. trichocarpa*. P value is calculated by one-sample *t*-test.

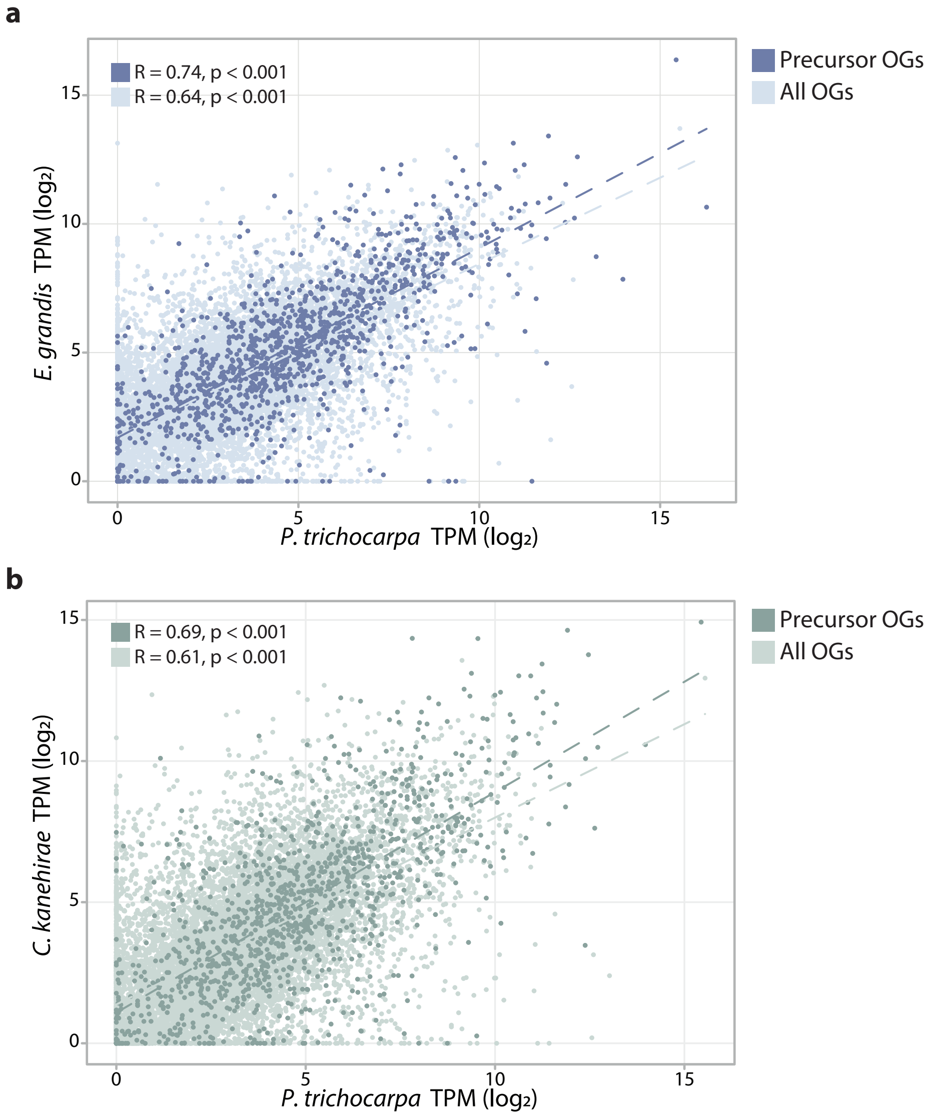

**Supplementary Figure 16. Sequence identity of sap peptide precursor regions between species.** Comparison of sequence identity (%) between *P. trichocarpa* and *C. kanehirae*, and between *P. trichocarpa* and *E. grandis*, focusing on sap precursor features. The box plots represent sequence identity in peptide-containing regions (orange) and non-peptide-containing regions (yellow) within the sap peptide precursor proteins. Asterisks denote statistically significant differences (Wilcoxon ranked sum test; $^{**}P < 0.01$)

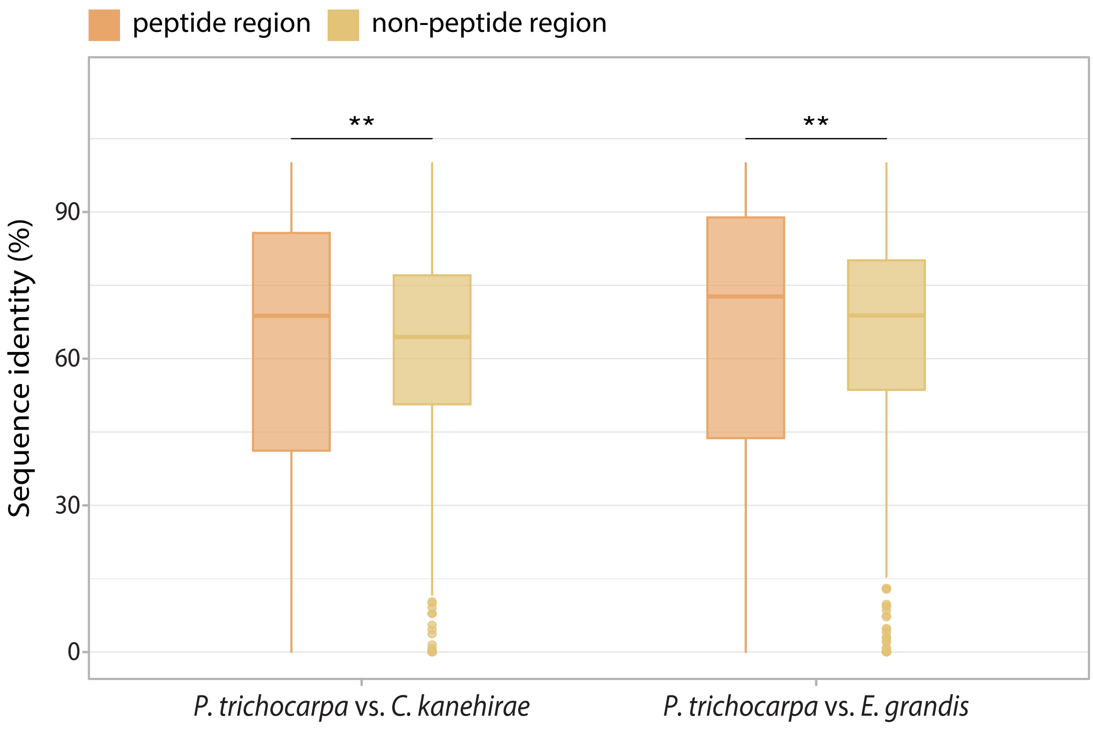

**Supplementary Figure 17. Comparative expression of sap peptide precursor gene orthologs between *P. trichocarpa*, *E. grandis*, and *C. kanehirae*.** Expression levels of sap peptide precursor gene orthologs identified from the *P. trichocarpa*, *E. grandis*, and *C. kanehirae* peptidome datasets are shown. Only orthologs with expression levels greater than 0.3% of their respective transcriptomes are illustrated. Colors represent orthology relationships as denoted in the legends. Proportional values for the contribution of each gene to the transcriptome are indicated in the figure.

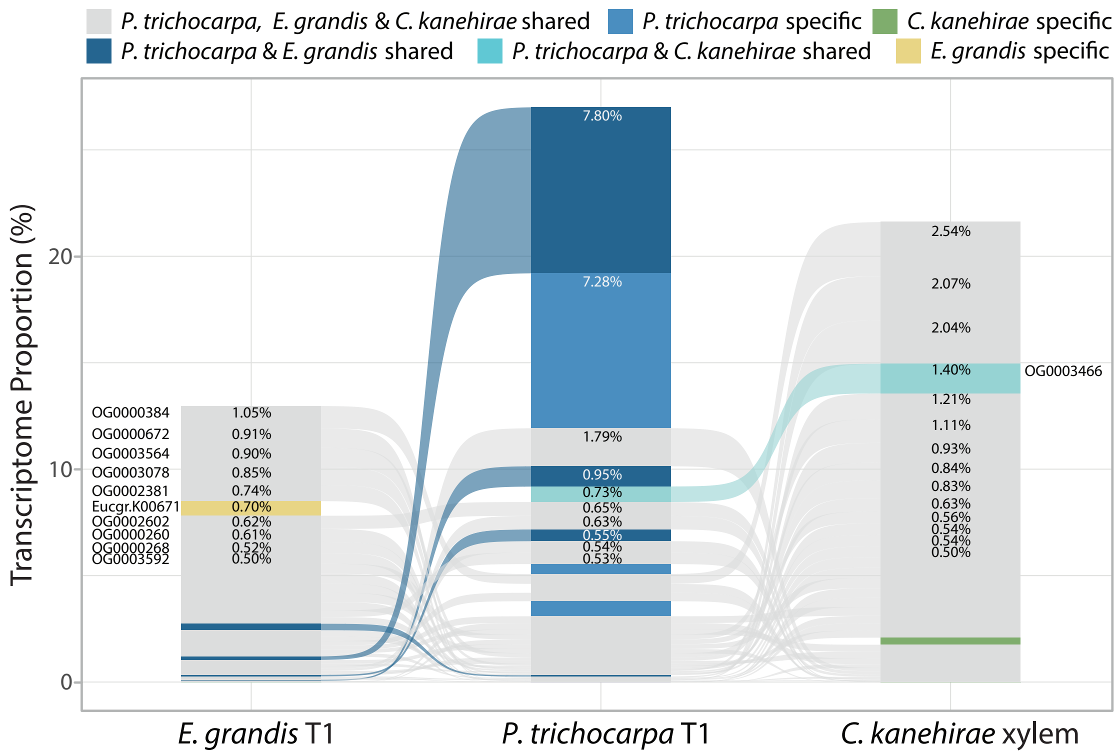

**Supplementary Figure 18. Overlap of orthogroups with peptide evidence in *P. trichocarpa*, *E. grandis*, and *C. kanehirae*.** Bars represent the orthogroup counts for unique and shared orthogroups, with connecting lines denoting combinations of orthology relationships across multiple species.

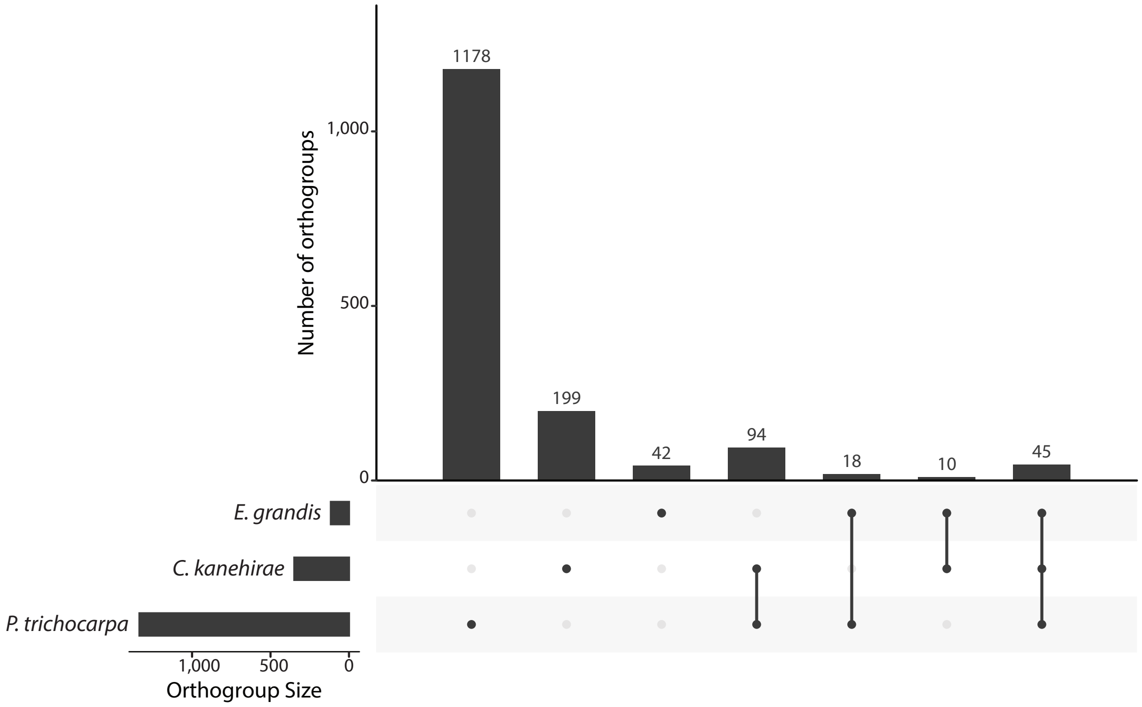

**Supplementary Figure 19. Proportion of unique peptide regions in *P. trichocarpa*.** The histogram shows the percentage of peptide regions unique to *P. trichocarpa* within orthogroups shared with *E. grandis* and *C. kanehirae*. Orthologous alignments were performed to identify distinct peptide regions supported by peptidome evidence. Only the most highly expressed orthologs containing sap peptides were used for comparison. Peptide regions in *P. trichocarpa* that did not overlap with those of other species were considered unique to *P. trichocarpa*. The proportion of these specific regions within each orthogroup was calculated.

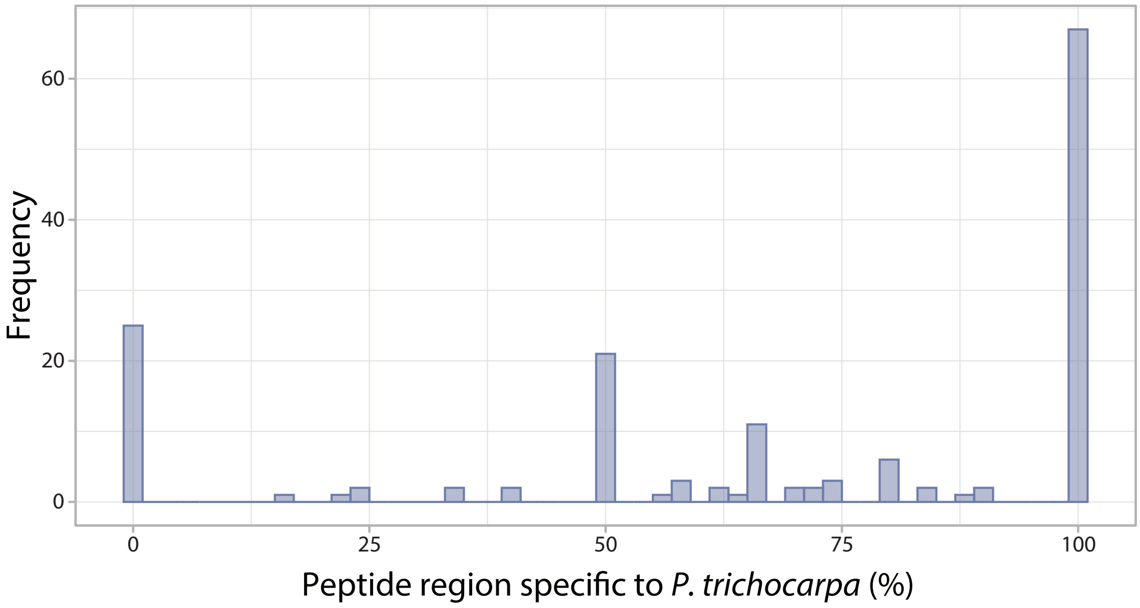
